# A Recombinant Approach For Stapled Peptide Discovery Yields Inhibitors of the RAD51 Recombinase

**DOI:** 10.1101/2023.02.24.529929

**Authors:** Teodors Pantelejevs, Pedro Zuazua-Villar, Oliwia Koczy, Andrew Counsell, Stephen J. Walsh, Naomi S. Robertson, David R. Spring, Jessica Downs, Marko Hyvönen

**Affiliations:** Department of Biochemistry, University of Cambridge, CB2 1GA, UK; Division of Cancer Biology, The Institute of Cancer Research, London SW3 6JB, UK; Yusuf Hamied Department of Chemistry, University of Cambridge, CB2 1EW, UK

**Keywords:** BRCA2, BRC repeat, macrocyclisation, peptides, stapled peptides, structural biology

## Abstract

Stapling is a macrocyclisation method that connects amino acid side chains of a peptide to improve its pharmacological properties. We describe an approach for stapled peptide preparation and biochemical evaluation that combines recombinant expression of fusion constructs of target peptides and cysteine-reactive divinyl-heteroaryl chemistry, as an alternative to solid-phase synthesis. We then employ this workflow to prepare and evaluate BRC-repeat-derived inhibitors of the RAD51 recombinase, showing that a diverse range of secondary structure elements in the BRC repeat can be stapled without compromising binding and function. Using X-ray crystallography, we elucidate the atomic-level features of the staple moieties. We then demonstrate that BRC-repeat-derived stapled peptides can disrupt RAD51 function in cells following ionising radiation treatment.

## Introduction

Peptide drugs contribute to 5% of the global pharmaceutical market, with the most notable example of recombinant insulin for diabetes treatment.^[1]^ Development of novel peptide-based drugs to address an ever-growing list of targets is hampered by a number of pharmacological pitfalls. Peptides suffer from short stability in biological fluids and low oral bioavailability, as they are prone to rapid proteolytic degradation.^[2]^ They are largely unable to cross the phospholipid membrane to engage intracellular targets. Macrocyclisation aims to improve some of these properties by constraining the conformation of a peptide.^[3],[4]^ This can render the peptide unable to fit into a protease active site, improving its stability and even oral bioavailability.^[5],[6]^ Macrocyclisation can also improve a peptide’s membrane permeability for intra-cellular targeting.^[7]^

Peptide stapling, in its broadest sense, is a macrocyclisation approach whereby the side chains of amino acid pairs within a peptide template are chemically linked to induce a constrained conformation.^[8]^ In the narrowest sense, stapling is the covalent linking of α-helical peptides using ruthenium catalysed metathesis (RCM) of non-natural amino acids bearing alkene side chains.^[9],[10]^ Recently, alternative structural elements, such as β-hairpins and loops have been successfully utilised for stapling.^[11],[12]^ Stapling of the nucleophilic side chain of cysteines has been explored as an attractive alternative to RCM, as it avoids the use of non-natural amino acids and metal catalysts.^[13]^ Stapling of cysteines can be done in mild, biocompatible conditions, allowing it to be used in the context of affinity selections of combinatorial libraries.^[14],[15]^

The stapling architecture, that is, the positions of the residues to be linked in the template, is a central variable in stapled peptide design. An appropriately placed staple should induce an unfavourable conformation or steric clash with the target. Stapling architectures are usually screened by preparing diverse peptide variants where the residue pair is systematically “scanned” along the sequence, or in a rational, structure-guided way when structural information is available. For helical peptides this is typically done by testing residues at fixed distance from each other, on the same side of an α-helix, while for other kinds of peptides in the absence of structural information screening is more complicated to design.

In this work, we explore the application of a recently developed class of bis-electrophilic divinyl-heteroaryl linkers for the recombinant production of cysteine-stapled peptides. Using the RAD51:BRC repeat interaction as a model template, we demonstrate that cysteine-stapled peptides can be prepared from small-scale bacterial cultures and screened for binding to evaluate different stapling architectures. The methodology provides an accessible, sustainable and rapid alternative to solidphase synthesis and allows stapling architectures to be evaluated in three days, starting from the initial cloning experiment. We then characterise the atomic-level structural changes in the binding modes of these peptides induced by stapling, showing that both helical and non-helical structural motifs can be linked with the divinyl-heteroaryl linkers. We then demonstrate that these peptides maintain functional activity in biochemical and cellular assays.

## Results and Discussion

### Small-scale Recombinant Preparation of Stapled Peptides

Solid-phase peptide synthesis (SPPS) of a linear precursor, followed by cyclisation and HPLC purification steps are typically performed to obtain stapled peptides for screening in biochemical or cellular assays.^[10],[16]^ Not only is this time-consuming and involves the use of harsh chemistry, certain peptides are difficult to synthesise and the process may require access to a peptide synthesiser. In order to rapidly evaluate different stapled peptide architectures, we set out to develop an alternative, small-scale strategy for peptide preparation and screening, using peptides produced in bacteria. This process allows for fast, quantitative determination of *in vitro* binding affinities (**Figure 1A**). The peptide is first designed by introducing a suitable pair of cysteines for linking in a linear template. Ideally, the introduction of cysteines is guided by an atomic structure of the template complexed with a target to identify mutable amino acid positions at suitable geometry and distance, but can also be performed in a structure-agnostic manner, for example, by predicting which residues are solvent-exposed and therefore not involved in binding.

**Figure 1.**
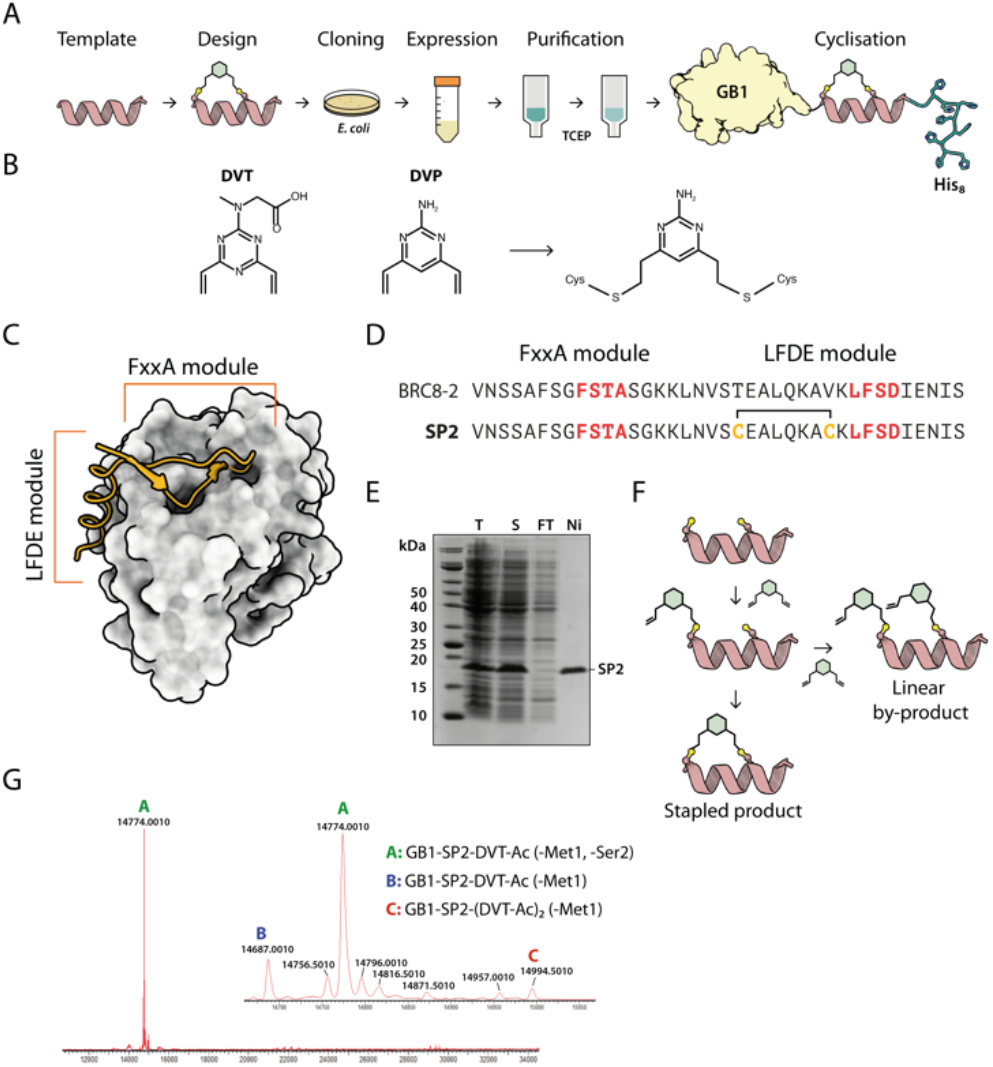
A recombinant approach for producing GB1-fused stapled peptides. (A) Work flow depicting the steps of stapled peptide preparation. The α-helix represents a generic template. (B) Divinyl-heteroaryl linkers used have been previously described.^[17],[18]^ (C) Structural model of a BRC repeat binding to RAD51 (PDB: 6HQU). (D) Sequences of the linear BRC8-2 template and **SP2**. (E) Coomassie-stained SDS-PAGE of eluted GB1-**SP2**-His_8_ model peptide (F) Mass spectrum (ESI-MS) of the optimised reaction products of GB1-**SP2**-His_8_ stapled with DVT. N-terminally excised residues are indicated in parentheses.

To facilitate this work, we have created a bacterial expression plasmid vector, pPEPT1, where the peptide is fused to an N-terminal Strep-Tag II, expression- and solubility-enhancing GB1 fusion partner (GB1) and a C-terminal octa-histidine tag (C-His_8_). The GB1 domain is relatively small and readily folded and expected to have minimal effect on the peptide fused to it. The C-terminal His_8_-tag is used for purification of the fusion protein, ensuring removal of possibly degraded peptide-fusions. The N-terminal StrepTag enables tandem-affinity purification should that be needed. DNA encoding for the peptide is assembled from synthetic oligonucleotides and cloned by sequence and ligation independent cloning (SLIC) which imposes no sequence constraints to the peptide or the linkers, including restriction enzyme recognition sites. The fusion protein carrying the peptide is expressed in a small scale (10 ml) *E. coli* culture, after which the cells are chemically lysed and loaded on an immobilized metal chelate affinity chromatography (IMAC) spin column. In order to prevent disulfide formation, the peptide cysteines are reduced on-resin using tris-carboxyethyl phosphine (TCEP). The peptide is eluted and directly reacted with a divinyl-heteroaryl linker in a double conjugate addition step, forming the staple moiety. The cysteine-selective linkers used in this work have been reported previously both in the context of peptide stapling (DVT, **Figure 1B**) and as tools for bio-conjugation of proteins (DVP, **Figure 1B**).^[17],[18]^ The linkers contain a six-membered core with two vinyl arms that yield a single stereoisomer of a stapled product. The linkers can also be decorated with other functional moieties, such as fluorophores.^[17]^ After the stapling reaction, excess linker is quenched by the addition of a thiol and the reaction products are used directly in a biochemical assay. The whole process from cloning to evaluation can be completed in three days and requires a basic biochemistry laboratory set-up, and can be performed in parallel with multiple peptide designs.

RAD51 is the central recombinase enzyme that catalyses mitotic homologous recombination, a key pathway of doublestrand DNA break repair.^[19]^ This process requires the assembly of an oligomeric RAD51 filament on resected ssDNA ends, mediated by an oligomerisation epitope located between its N- and C-terminal domains.^[20]^ RAD51 is regulated by a set of conserved ~35 aa long BRC repeats located in the BRCA2 tumour suppressor protein.^[21,22]^ BRC repeats bind the C-terminal ATPase domain of RAD51 via two conserved tetrad motifs, FxxA and LFDE, located on eponymous sequence modules (**Figure 1C,D**).^[21]^ The FxxA motif is also found on the RAD51 oligomerisation epitope, leading to competition with the BRC repeats for the same interface. We used the RAD51:BRC repeat interaction as a model system to evaluate our stapling methodology because the long BRC peptide permits a large number of different stapling architectures to be tested in a single template. As template, we chose a previously identified high-affinity shuffled repeat BRC8-2, which binds a monomeric construct of RAD51 (HumRadA22) with a low-nanomolar K_D_ and inhibits RAD51 oligomerisation on ssDNA *in vitro*, and for which a complex structure has been determined.^[23]^ Model peptide **SP2** was designed by introducing two cysteines at the α-helical, C-terminal LFDE module of the 38-residue template (**Figure 1D**). The designed cysteines replaced solvent-exposed residues *i*, *i*+*7* positions apart in a classical helical stapling fashion. Small-scale expression yielded the linear peptide fusion on a 10 μmol scale and its purity was confirmed by SDS-PAGE (**Figure 1E**).

We proceeded with optimising the stapling reaction of the linear **SP2** precursor. Successful cysteine stapling involves a two-step mechanism (**Figure 1F**). The first step is a bimolecular reaction between the nucleophilic cysteine and an electrophilic arm of a linker, followed by an intramolecular ring-closing reaction of the second arm with the second cysteine. The second ringclosing step can be in competition with an undesired bimolecular side-reaction, where an additional linker molecule reacts with the second cysteine. The resulting linear double-linker product is of no pharmacological utility. Typically, such side-products can be separated by HPLC, however, our method is specifically aimed at avoiding the use of specialist chromatography equipment. In order to slow down the rate of the unwanted second linker addition and ensure the formation of the correct product, we used pseudo-dilution of the linker via its step-wise addition to the linear **SP2** peptide. To optimise the reaction, we examined a number of conditions, such as the rate at which the linker is added to the peptide, as well as pH and presence of TCEP. Using mass spectrometry (ESI-MS), we observed that the majority product of the initially trialled stapling reaction is the correctly linked cyclic peptide (**Figure S2**, reaction **a**; **Figure S4**). The double linker side-product was also observed at a much lower peak intensity and its abundance correlated with the pace of linker addition, confirming that pseudo-dilution can aid quantitative cyclisation (**Figure S2**, reactions **a,b,c; Figure S4-S5**). We assessed if TCEP can be included in the reaction to ensure cysteines are reduced and to minimise the formation of disulfides. However, we found that at 500 μM TCEP rapidly reacts with the linker, forming undesired linear peptide-linker-TCEP adducts (**Figure S2**, reactions **d,e,f; Figure S7-S9**). Considering this, peptides were subsequently reduced on-resin and the stapling reaction performed immediately after elution. The final optimised reaction yielded a highly pure cyclised peptide fused to an N-terminal GB1 tag and a C-terminal His_8_-tag, as evidenced by ESI-MS (**Figure 1F, Figure S2**, reaction **h**).

### Affinity screening of stapled peptides

Having established a procedure for producing recombinant, cysteine-stapled peptides, we set out to prepare a variety of designs and evaluate these in a binding assay. Peptides were designed in a structure-guided manner, informed by the previously published BRC8-2:RAD51 complex structure (PDB: 6HQU).^[23]^ Spatial proximity and residue geometry were used as criteria for cysteine placement. Facile modelling suggests that inter-sulphur distances of 4-10 Å may be suitable for stapling with the divinyl-heteroaryl linkers. Previously it was shown that a α-helical peptide can be stapled with the DVT linker at *i*,*i*+*7* positions without perturbing its secondary structured.^[18]^ In a traditional α-helical stapling approach, we introduced different *i*,*i*+*7* cysteine pairs at the C-terminal LFDE module of the BRC8-2 repeat: **SP1**, **SP8**, **SP9, SP15** in addition to the model peptide **SP2** (**Figure 2A and Table 1)**. Residues selected for mutagenesis were solvent-exposed and did not form any apparent interactions with the protein. Alternatively, peptides were designed with at least one cysteine located at the β-hairpin-containing FxxA module (**Figure 2A and Table 1; SP10**, **SP11, SP12**, **SP13**, **SP14**, **SP16**).

**Figure 2.**
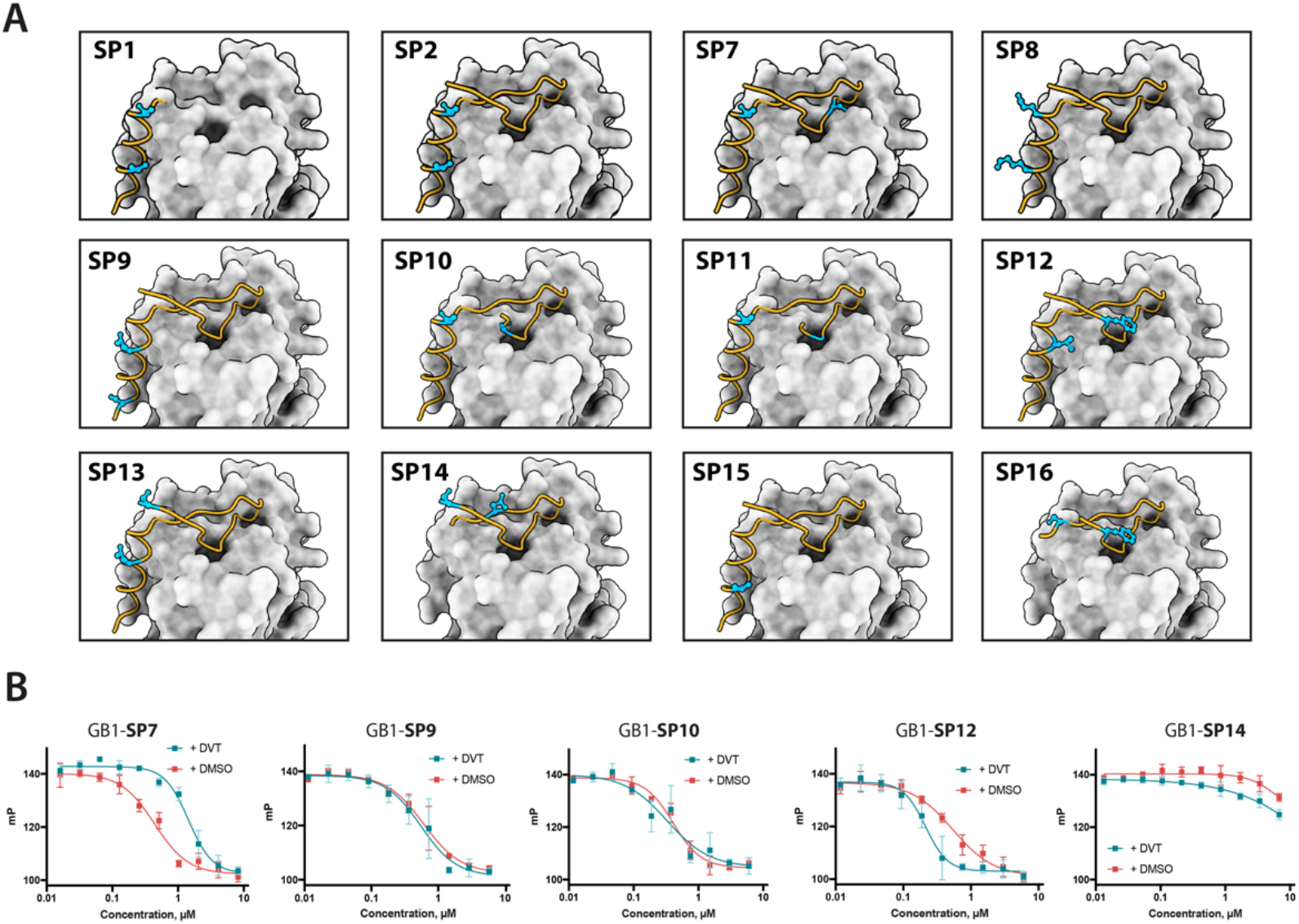
Design and screening of stapled BRC8-2 peptides. (A) Structural model of BRC8-2 peptide bound to HumRadA22 with mutable residues highlighted and side chains visualised for each design. SP15 lacks the second mutable residue as it is not defined in the complex structure. (B) Representative FP assay measurements. Red curves are unstapled, linear controls; green curves are titrations of stapled peptides. Data shown is the mean of three replicates ± SE. Data was fitted using a four-parameter logistic equation.

**Table 1.**
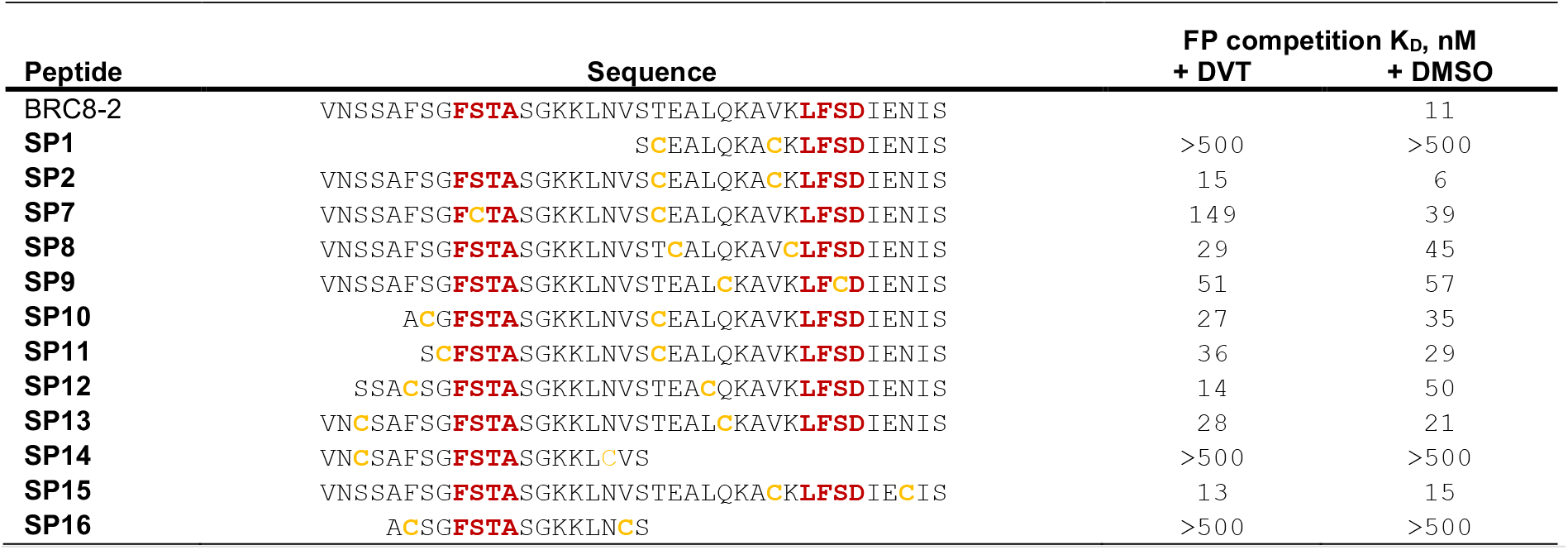
Sequences of stapled BRC8-2 peptides and their stapling architecture, as well as the KD values determined from the FP competition assay. KD values were calculated from IC50 values using a previously reported equation.^[26]^ Titrations were done as single experiments with three technical replicates at each titration data point.

Placement of cysteines in these designs was likewise guided by the structure of the complex. For example, **SP12** mutates a solvent-exposed residue Phe2055 near the N-terminus of the β-hairpin and a Leu1234 at the middle of the α-helix of the LFDE module. Both side chains are solvent-exposed and located nearby (d_Cα-Cα_ = 11 A), despite an 18 aa sequence distance. A number of peptides (**SP1, SP14, SP16**) contained complete truncations of either the FxxA or the LFDE module. We also included a negative control peptide (**SP7**), where the stapled residues are so far from each other that their stapling is expected to disrupt the binding mode, rendering the peptide unable to maintain the FxxA and LFDE hot-spot interactions with RAD51 simultaneously.

We prepared all of the stapled peptides from small-scale bacterial cultures as GB1/C-His_8_ fusions and cyclised them with the DVT linker. For peptides **SP10**, **SP11**, **SP12** and **SP15** correct stapling was confirmed by MS, showing a similar composition to **SP2** (Supplementary Data: Mass spectra of representative GB1-fused stapled peptides, **Figure S12-S15**). Thus, we confirm that the pseudo-dilution approach is robust in yielding almost exclusively the cyclised product, irrespective of whether a helical or non-helical motif is being constrained. To examine the effect of introducing a covalent staple on affinity, a mock reaction was done in parallel for each peptide by splitting the IMAC elution into two halves and adding DMSO to the mock control instead of the linker. TCEP (500 μM) was included in the mock reaction to maintain the peptide in linear form by preventing disulfide formation. The peptides were tested in a fluorescence polarisation (FP) assay monitoring the displacement of a fluorescently-labelled BRC4 repeat from a monomeric version of RAD51 (HumRadA22), as reported previously.^[23]^ Fitted K_D_ values and representative dose-response curves are provided in **Table 1** and **Figure 2B**. The negative control peptide **SP7** had a K_D_ of 39 nM after mock stapling, whereas the stapled product has a K_D_ of 149 nM, a more than a three-fold reduction in affinity, confirming the disruption of binding by an inappropriately introduced staple. All of the peptides containing both FxxA and LFDE hot-spot motifs bound HumRadA22 with high affinity after stapling (K_D_ of 15-51 nM). For these peptides, we observed minimal differences in affinity between the corresponding stapled and mock forms.

Because the experiments were conducted as single titrations of technical triplicates at each concentration, we do not compare the K_D_ values of these high-affinity peptides. We did not observe high-affinity binding for any of the significantly truncated peptides, expanding the previous observation that both the FxxA and LFDE motifs are critical for the interaction of the BRC4 repeat.^[24],[25]^ None of the repeats, either stapled or linear, bound with a higher affinity than the BRC8-2 template (K_D_ = 11 nM).

### Biochemical characterisation of cysteine-stapled BRC8-2 repeats

To evaluate functional and structural properties of stapled BRC repeats, we prepared peptides in a tag-free form. For this, we expressed them in scaled-up *E. coli* cultures as fusions to an N-terminal His-tag, followed by a GB1 fusion and a TEV cleavage site. Peptides were purified by IMAC, cleaved proteolytically, stapled using the pseudo-dilution approach in scaled-up reactions, and purified by HPLC, which yielded each peptide in >5 mg yield. Because both the GB1 and His-tag are N-terminal, the final products contained just the stapled peptide with a 1 to 2-residue linker at the N-terminus to ensure efficient cleavage by the TEV protease, as confirmed by LCMS (see **Figure S16-19**). Three stapled BRC8-2 repeats were produced in this manner (**SP2, SP24, SP30**). **SP2** and **SP24** contain cysteines at identical positions in the LFDE module but differ in length and sequence. **SP30** is based on **SP12** and contains a distant *i*,*i*+18 linkage across the two modules. **SP24** and **SP30** were significantly truncated compared to the template to remove residues that are unlikely to contribute to binding, as suggested by the BRC8-2:RAD51 structure. Truncation of flexible termini can aid crystallisation of a peptide:protein complex and may also be beneficial for cellular uptake. To prevent a double negative charge at the C-terminus arising from the truncation, the resulting C-terminal Asp in **SP24** and **SP30** was mutated to Gly. A C-terminal Gly results in a flexible acidic tail that can mimic the Asp side chain. A different linker (DVP, **Figure 1B**) was used to cyclise **SP24** and **SP30**, instead of DVT which was used for **SP2**. DVP lacks a negative charge and may benefit cellular uptake.

Previous studies have demonstrated that stapling can preorganise unbound peptides towards the bound conformation.^[18],[27],[28]^ In particular, rationally introduced staples were shown to enforce α-helices. This effect has been hypothesised to be responsible for the improved membrane permeability of stapled peptides.^[29]^ We characterised the secondary structure content of the three peptides **SP24** and **SP30** by circular dichroism (CD), comparing their stapled and linear versions with the BRC8-2 template (**Figure 3B**). All of the peptides had a mixed secondary structure composition, with a global minimum at 200 nm corresponding to a random coil and a smaller minimum at 225 nm suggestive of an α-helix. Stapling of **SP24** increases its α-helical character, evidenced by a more negative molar ellipticity in the 210-230 nm range. This is consistent with stabilisation of the LFDE α-helix by the staple. No significant change in secondary structure was observed for **SP30**, indicating that the large cycle likely maintains a high degree of flexibility.

**Figure 3.**
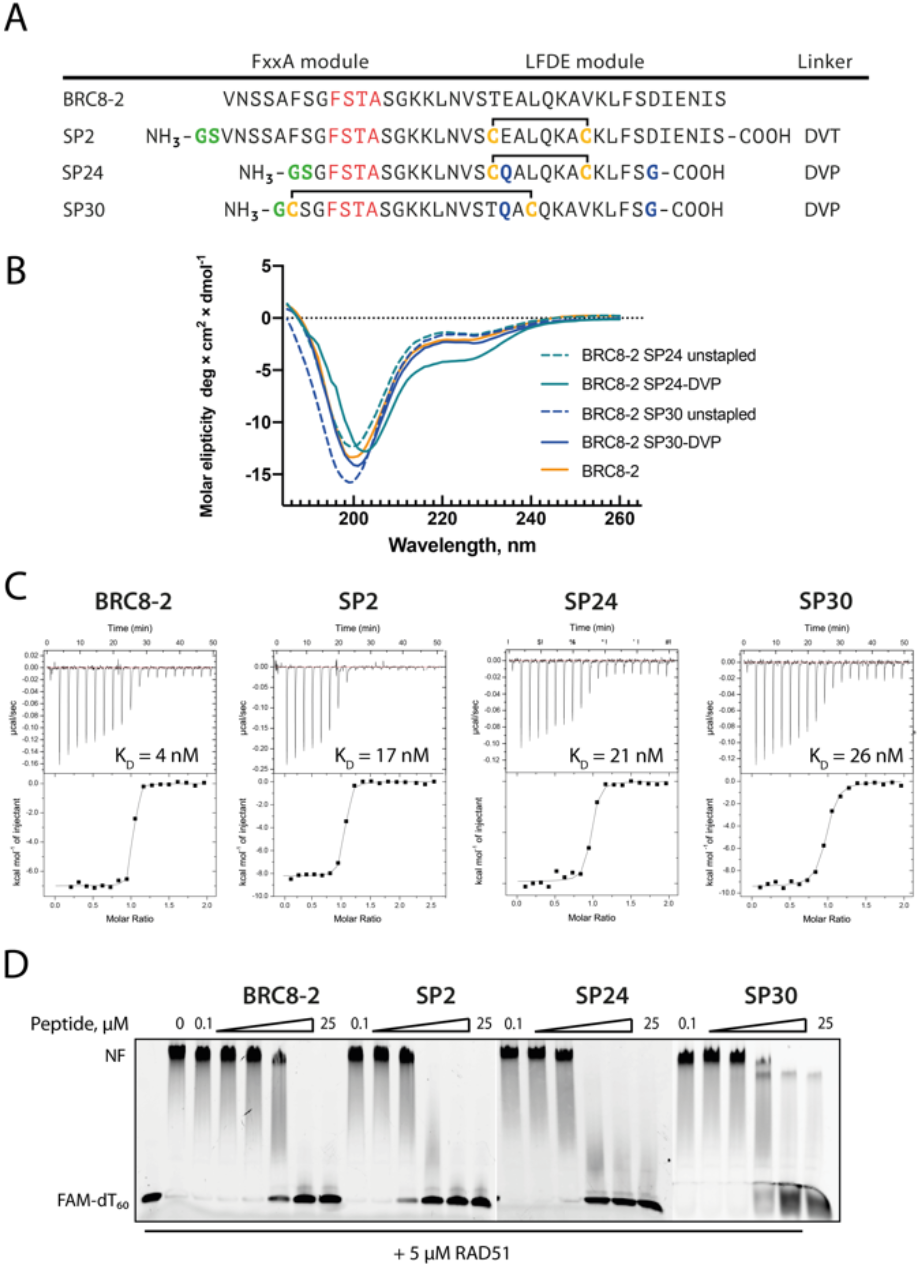
Characterisation of stapled BRC8-2 repeats purified as free peptides. (**A**) Sequences of **SP2, SP24, SP30,** as well as the linear template. (**B**) Circular dichroism spectra of **SP24** and **SP30** in stapled and reduced linear forms. (**C**) ITC measurements of peptide binding to the HumRadA22 protein. (**D**) EMSA of RAD51:dT_60_ nucleoprotein filament incubated with varying concentrations of peptides.

We then evaluated the binding of **SP2**, **SP24** and **SP30** to HumRadA22 using isothermal titration calorimetry (ITC, **Figure 3C)**. All three peptides bound with similar low-nM affinities, and, similarly to the FP measurements with the GB1-fused constructs, binding appears to be slightly weaker compared to the linear BRC8-2 template.

Isolated BRC repeats disrupt the RAD51-ssDNA nucleoprotein filament by competing with RAD51 oligomerisation interface.^[30,31]^ We evaluated the effect of stapled peptides **SP2**, **SP24** and **SP30** on the nucleoprotein filament using electrophoretic mobility shift assay (EMSA, **Figure 3D**). A fluorescent dT_60_ ssDNA was incubated with full-length RAD51, after which peptides were added at different concentrations before separating the products on a polyacrylamide gel. All three peptides depolymerise RAD51 from ssDNA to similar extent, suggesting that the peptides maintain function in the context of the full-length human protein.

### Structural characterisation of cysteine-stapled BRC8-2 repeats

Atomic-level characterisation of peptide:target complexes has revealed the impact of a variety of stapling chemistries on the binding modes of stapled peptides.^[11],[32],[33],[34]^ Understanding the conformational changes induced by the staple moiety can benefit the design of new binders. While it has been shown that DVT can link *i*, *i*+*7* residues on α-helical MDM2-binding peptides, the conformation of the linker and its effect on the helix geometry are unclear. To elucidate their binding modes, we co-crystallised **SP2**, **SP24** and **SP30** in complex with HumRadA22. Crystals of all three complexes yielded high-resolution structures, revealing the atomic-level detail of the staple. In the lowest resolution **SP2** complex structure, the DVT linker appears as a poorly defined “blob” in the electron density located mid-way between the two cysteine sulphurs (**Figure 4A**). In the **SP24** complex structure, the DVP linker can be observed in atomic detail and its conformation can be modelled (**Figure 4B**). The two arms of the DVP linker in **SP24** have acquired different orientations relative to the heterocycle. The first arm, connected to Cys1231, has the C-C bond perpendicular to the pyrimidine ring plane, whereas in the arm linking Cys1238, the bond is nearly co-planar. In both arms, the C-C bond and the C-S bonds resemble a *trans* conformation. Thus, the staple moiety appears to experience little or no torsional and steric strain to accommodate the *i*, *i* + *7* α-helical link.

**Figure 4.**
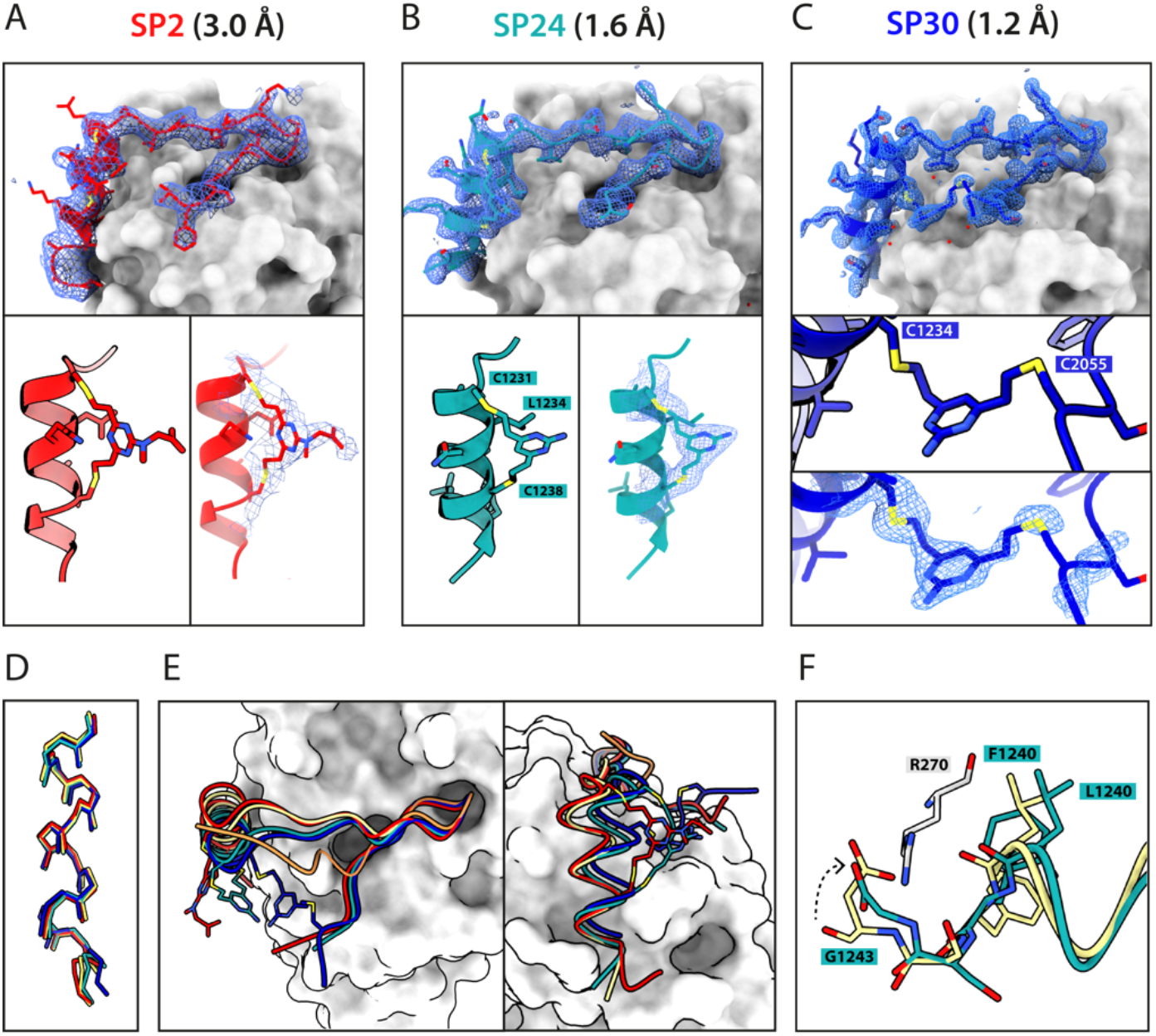
Crystal structures of peptide complexes. Mesh depicts weighted 2mFo-DFc electron density maps after several rounds of refinement. Binding modes and zoomed-in staple moieties are shown for (A) **SP2**, (B) **SP24** and (C) **SP30**. (D) Alignment of the α-helical LFDE modules from the three stapled peptides and the BRC8-2 template complex (tan). (E) Backbone movement relative to the template structure observed for **SP24** (green) and **SP30** (blue). Complexes were superposed based on the HumRadA22 Cα atoms. (F) Rotation of the LFDE α-helix in **SP24** results in a shift of Leu1240 and Phe1241 side chains relative to BRC8-2 template. Gly1243 forms a salt bridge with HumRadA22 Arg270 (*Hs*RAD51 Arg254), mimicking an acidic sidechain.

The linker in **SP24** does not interact with HumRadA22, but its heterocycle stacks on the Leu1234 side chain, creating a hydrophobic cluster. In both **SP2** and **SP24**, the linker lies in an unoccupied region between the side chains of endocyclic *n*+*3* and *n*+*4* residues. The α-helical geometry of the LFDE module is not perturbed either in **SP2** or **SP24**, with near-identical distances observed between cysteine α-carbons compared to the corresponding residues in BRC8-2 (**Figure 4D**). The Cα RMSD of the superimposed helices from **SP2** and **SP24** relative to BRC8-2 are 0.235 and 0.310 Å, respectively, indicating minimal distortion by *i*,*i*+*7* stapling. These structural observations support the application of divinyl-heteroaryl linkers as a general strategy for *i*,*i*+*7* stapling of α-helical epitopes.

In the **SP30** complex, the linker connects Cys2055 and Cys1234, creating a 19 aa cycle (**Figure 4C**). Electron density is clearly defined for the side chain and the linker arm on the Cys1234 side, whereas the other arm is less interpretable, possibly due to conformational flexibility. The pyrimidine core makes contacts with HumRadA22 residues Gln213 and Gln217. Compared to the helical staples in **SP2** and **SP24**, the linker arms in **SP30** acquire different conformations, where both C-C bonds are perpendicular to the heterocycle core.

The C-C bond at Cys1234 and C-S bond at Cys2055 are in a *gauche*-like conformation, suggesting that unfavourable local strain accommodates the binding mode.

All three stapled peptides have broadly similar binding modes, with FxxA and LFDE hot-spot residues binding to cognate hydrophobic pockets. However, the three stapled peptides lack the extended β-hairpin observed for BRC8-2. Instead, the N-terminal residues Ser2052-Gly2057 point away from the peptide and do not contribute to intra-molecular H-bonding. This is anticipated in **SP30**, where this region is perturbed by the introduction of a staple, however, it is not clear why **SP2** and **SP24** display such change. It is possible that the N-terminal movement is induced by crystal contacts and not as a result of stapling. Remarkably, the LFDE modules of **SP24** and **SP30** undergo substantial movement relative to HumRadA22 and the rest of the peptide (**Figure 4E**). The motion affects more than half of the peptide, encompassing residues Lys1226 to Glu1243. This is concomitant with a rotation of the LFDE α-helix around its helical axis, leading to the re-organisation of Leu1240 and Phe1241 side chains in their cognate pockets (**Figure 4F**). It is reasonable to hypothesise that the shift of the LFDE module is a consequence of the described C-terminal Asp1243Gly mutation introduced to reduce the overall negative charge of the peptide. The carboxylate of the terminal glycine residue is one carbon shorter compared to an aspartate side chain, and rotation of the LFDE helix brings it closer to HumRadA22 Arg270 (*Hs*RAD51 Arg254) in the **SP24** structure, in order to maintain the salt bridge. Despite this large-scale shift, the peptides maintain low nanomolar affinities, suggesting that the RAD51:BRC interface can accommodate structural plasticity not observed in previous studies.

### Cysteine-stapled BRC repeat inhibits RAD51 foci formation in cells

With some exceptions, the large size of peptides is detrimental to membrane permeability and restricts their use to extra-cellular targets. Various cationic cell-penetrating motifs have been conjugated to peptides and other biomolecules to aid cellular uptake. Arginine-rich cell-penetrating peptides (CPPs) are the most commonly applied motif that has been shown to internalise by inducing membrane multilamerality and forming of a fusion pore.^[35]^ To enhance cellular uptake of our stapled BRC repeats, we prepared an polyarginine derivative of **SP30**, called **SP31**, by recombinantly introducing and N-terminal Arg_9_ sequence (**Figure 5A**).

**Figure 5.**
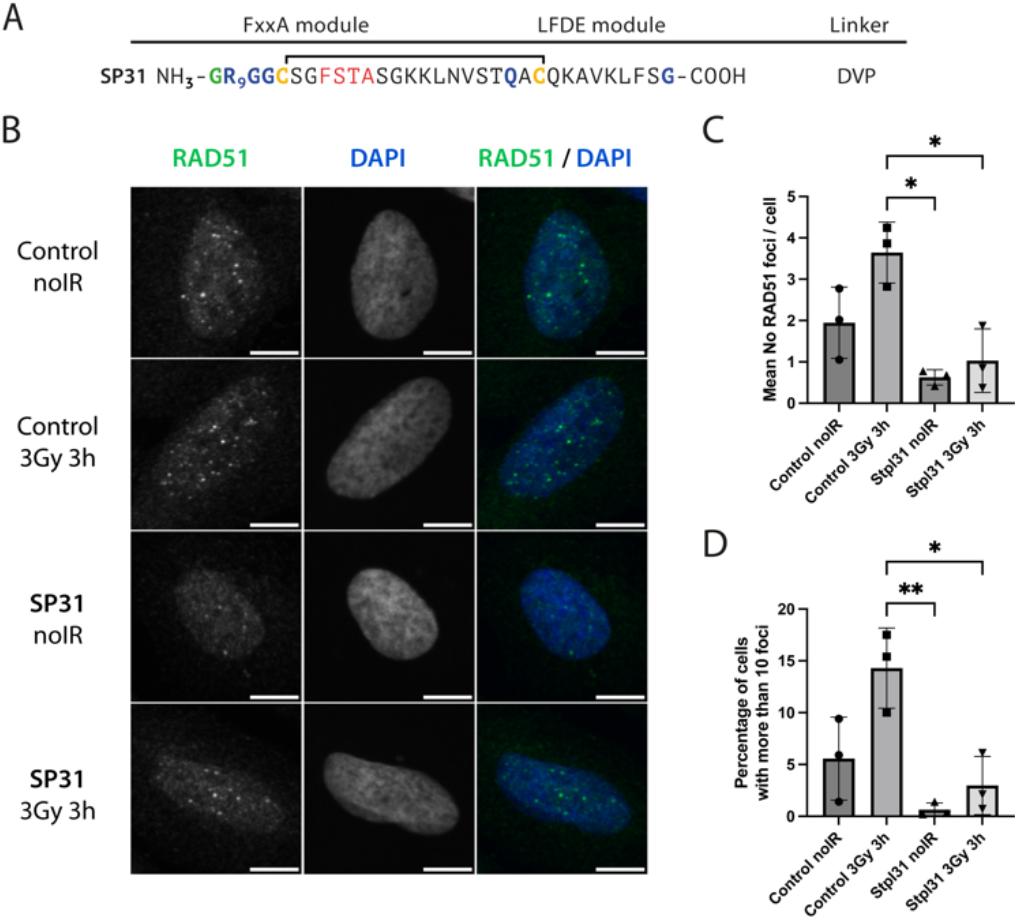
**SP31** disrupts RAD51 foci formation in U2OS cells. (A) Sequence of **SP31** (B) Representative IF images of U2OS cells incubated with **SP31** (40 μM, SP31) or vehicle alone (Control) for 1 hour, after which they were treated with 3Gy ionising radiation (3Gy 3h) or no radiation (noIR) and recovered for 3 hours. Cells were stained with α-RAD51 and DAPI as indicated. Scale bar is 10 μm. (C) Bar graph showing the average of the mean counts of RAD51 foci per cell from three independent biological experiments. Data are presented as mean ± SEM, * - P < 0.05, using ANOVA test followed by Tukey’s method. (D) Bar graph showing the average percentage of U2OS cells with more than 10 foci per cell. Data are presented as mean ± SEM, * - P < 0.05, **- P < 0.01, using ANOVA test followed by Tukey’s method.

Following treatment of cells with ionising radiation (IR), RAD51 translocates to the nucleus and forms foci at sites of DNA damage that are visible by immunofluorescence (IF) microscopy. Previously, we have shown that transient expression of a GFP-BRC8-2 peptide fusion leads to the efficient disruption of IR-induced RAD51 foci in the U2OS osteosarcoma cell line.^[23]^ We monitored the ability of **SP31** to disrupt IR-induced RAD51 foci formation using IF. Cells were pre-incubated with **SP31** or with vehicle (control), after which they were either treated with no radiation or with 3Gy IR and imaged after a 3-hour recovery. We observed RAD51 foci in the absence of peptide and IR treatment, which likely reflects recombination events associated with the basal replicative stress of U2OS cells (**Figure 5B-D, Figure S3**). As expected, IR treatment of control cells leads to an increase in the number of RAD51 foci, however this does not reach statistical significance under the experimental conditions used (**Figure 5C-D, Figure S3)**. Notably, addition of **SP31** reduces the number of RAD51 foci in both IR-treated and untreated cells, with a more than three-fold decrease in mean foci counts in the IR-treated sample (**Figure 5B-D, Figure S3**). The IF data demonstrates that the Arg9-fused peptide **SP31** is able to pass cell membrane and sequester RAD51, preventing its localisation to DSB sites and oligomerisation on DNA.

## Conclusion

Finding an optimal stapling architecture requires the testing of many peptide sequences. Preparation of stapled peptides is traditionally performed using specialist equipment such as peptide synthesisers, lyophilisers and HPLCs and uses toxic solvents such as dimethylformamide (DMF) and N-Methyl-2-pyrrolidone (NMP). This renders solid-phase peptide synthesis challenging from a sustainability standpoint, calling for specific waste-disposal procedures. Procurement of a large number of synthetic peptides can be cost-prohibitive, with typical prices at $10 USD per amino acid for 90% pure product. We have presented a workflow that allows stapled peptides to be prepared in sufficient amounts for quantitative evaluation of their binding using standard biochemistry equipment. Utilising parallel, smallscale bacterial expression of peptides as small fusion proteins with tags for rapid purification in combination with cysteine-reactive linkers, we demonstrated that stapled peptides can be prepared in three days from the initial cloning steps. The insert encoding for the peptide is generated with accessible and inexpensive oligonucleotides; a pair of oligos needed for a 30-residue peptide costs ca. $20 USD. The GB1 fusion improves expression levels and increases peptide solubility, and allows for quantification by UV absorbance. This approach will be suited for proof-of-concept studies with the aim of modulating proteinprotein interactions. However, compared to use of purified synthetic peptides, some limitations warrant discussion as well. In some contexts, the GB1 fusion may exert a detrimental effect on binding due to steric repulsion with the target protein and we advise to introduce a longer amino acid linker to avoid this. Other fusion partners could also be used, provided they do not contain cysteines, and our pPEPT1 vector is built so that the GB1 can be replaced easily with, for example, SUMO (small ubiquitin-like modifier) protein. Secondly, the method involves a single purification step with the C-terminal His-tag, therefore proteolysis products may be co-purified and interfere with downstream assays. To counter this, we have included in the plasmid an N-terminal StrepTag II for tandem affinity purification.

The small-scale preparation of GB1-BRC8-2 stapled peptides yielded sufficient product for testing in a FP assay. We envisage that peptides prepared in this manner can be used in a range of assays, such as surface plasmon resonance (SPR), homogeneous time-resolved fluorescence (HTRF), dynamic scanning fluorimetry (DSF) and others. The methodology is not suitable for cell assays where the target is intracellular, as the GB1 tag is most certainly preventing cellular uptake. Instead it is aimed at cases where a purified target is available for *in vitro* measurements. We anticipate that the described method can be used with alternative Cys-reactive linkers, such as bis-haloacetamides, however, reaction selectivity will have to be optimised and it is not certain that similar levels of stapled product can be obtained.

We have used the BRC8-2:RAD51 complex structure to inform the design of a structurally diverse set of stapled peptide variants. Using our recombinant workflow, we show that different structural motifs in the BRC repeat can be stapled without impairing affinity. We also show that BRC repeats can be truncated with no disruption in binding. However, both FxxA and LFDE motifs are indispensable, supporting previous studies. Our crystallographic analysis reveals the different structural features of the staple, which will be beneficial for future studies utilising these linkers. Finally, these peptides potently de-polymerise RAD51 from ssDNA *in vitro* and, when fused to a cell-penetrating peptide, inhibit RAD51 foci formation in cells. The stapled peptides described in this work represent a novel modality for targeting RAD51 by competing with its self-oligomerisation and interaction with BRCA2.

## Acknowledgements

We are grateful for Dr Laurens Lindenburg for the gift of BRC4-fluorescein. We thank the Biophysical and X-ray crystallographic facilities (Department of Biochemistry, University of Cambridge) for access to instrumentation and support. We thank Diamond Light Source for access to beamlines i03 and i04 (proposals mx18548, mx25402). TP was supported by MRC DTP. The Spring Laboratory acknowledges support from the EPSRC, BBSRC and MRC. PZ-V and JAD were supported by CRUK (C7905/A25715).

