## Supporting Information for "A Recombinant Approach For Stapled Peptide Discovery Yields Inhibitors of the RAD51 Recombinase"

[4] Present address: Latvian Institute of Organic Synthesis, Aizkraukles 21, Riga, LV-1006, Latvia

### Materials and Methods

#### Materials

| Reagent | Source | Identifier |
|---|---|---|
| **Cell lines** | | |
| U2OS | ATCC | HTB-96 |
| **Antibodies** | | |
| Rabbit anti-RAD51 | Santa Cruz biotechnology | sc-8349 |
| Mouse anti-rabbit Alexa Fluor 488 | Life Technologies | A21206 |
| **Bacterial strains** | | |
| *E. coli* T7 Express chemicompetent cells | Prepared in-house from a New England Biolabs glycerol stock | C2566H |
| *E. coli* BL21 (DE3) Rosetta | Prepared in-house from a Novagen stock | 70954-3 |
| **Recombinant DNA and oligonucleotides** | | |
| pPEPT1 plasmid | This work. See **Figure S1.** | Adgene #195001 |
| pOP3BT | Hyvonen Lab, University of Cambridge | Addgene #112603 |
| pEXP-NHis-GB1 | Hyvonen Lab, University of Cambridge | Addgene #112565 |
| Oligonucleoides for cloning peptides. | Synthesised by Sigma-Aldrich. See **Table S2**. | |
| **Purified proteins and peptides** | | |
| GB1-peptide fusions | This work | |
| Un-tagged BRC repeat peptides | This work | |
| Fluorescein-BRC4 | Gift from Dr Laurens Lindenburg, previously reported in [1] | |
| Human RAD51 | This work | |
| HumRadA22 protein | This work, previously described in [2] | |
| Lysozyme | Sigma-Aldrich | 1.05281 |
| T4 DNA polymerase | New England Biolabs | M0203S |
| DNAse I | Sigma-Aldrich | DN25 |

| | | |
|---|---|---|
| TEV protease | Prepared in-house using bacterial expression from the pRK793 plasmid (Addgene #8827) | |
| Phusion DNA polymerase | New England Biolabs | M0530S |
| *Bsa*I | New England Biolabs | R3733S |
| *Hind*III | New England Biolabs | R0104S |
| *Xho*I | New England Biolabs | R0146S |
| *BamH*I | New England Biolabs | R3136S |
| **Chemicals** | | |
| DVP linker | Spring lab, University of Cambridge, described previously[3] | |
| DVT linker | Spring lab, University of Cambridge, described previously[4] | |
| TCEP | Melford | T26500 |
| IPTG | Melford | I56000 |
| Ampicillin | Melford | A40040 |
| AEBSF | Melford | A20010 |
| Ni-NTA agarose | Cube Biotech | 31103 |
| NEBuffer 2.1 | New England Biolabs | B6002S |
| 2xYT medium | Formedium | AIM2YT0210 |
| Penicillin/streptomycin | Sigma-Aldrich | P4333 |
| Bovine serum albumin | Sigma-Aldrich | A9647 |
| Fetal bovine serum | Sigma-Aldrich | F7524 |
| DMEM – high glucose | Sigma-Aldrich | D6429 |
| Triton X-100 | Sigma-Aldrich | T8787 |
| ProLong Gold Antifade Mountant with DAPI | ThermoFisher | P36935 |
| **Columns and consumables** | | |
| HiTrap Heparin 5 ml | Cytiva | 17040703 |
| PD-10 desalting column | Cytiva | 17085101 |
| 384-well flat-bottom microplates | Corning | 3821 |

**Small-scale preparation of GB1-fused stapled peptides**

Peptide-coding DNA was designed using the DNAWorks online application,[5] optimising codon usage for *E. coli* expression. DNA oligonucleotides for the assembly of these fragments were generated using the same software and 15-20 nt linkers were appended to the 5' ends of the outermost forward and reverse oligos for sequence and ligation independent cloning (SLIC). Oligonucleotides used for assembly of the fragments are shown in Supporting Information: *Oligonucleotides for cloning of BRC repeat-coding DNA inserts*. The inserts were then synthesised by assembly PCR using Phusion DNA polymerase (New England Biolabs) with standard reaction conditions. Each assembly PCR reaction contained 1 μM of the outermost oligos and 0.02 μM of each internal oligo. Inserts were then purified by gel extraction using the GeneJet gel extraction kit (ThermoScientific). Sequence and ligation-independent cloning (SLIC) was used to clone the inserts into the pPEPT1 vector (Addgene #195001, **Figure S1**) digested with the *Bsa*I restriction enzyme. Using *BsaI* allows the peptide to be cloned in a seamless fashion as cleaves outside of its recognition site. 10 µl SLIC reactions contained purified insert at 2-10 ng/µl and digested vector at 4-20 ng/µl in 1x NEB Buffer 2.1 (New England Biolabs). 0.6 U of T4 DNA polymerase (New England Biolabs, M0203S) was added to each reaction and incubated for 1-5 min at RT, after which either dGTP or dCTP was added to a final concentration of 10 mM to stop exonuclease activity. Reaction was further incubated for 1 minute at RT, after which it was heated for 5 minutes at 65 °C in a PCR cycler to deactivate the T4 polymerase, after which the PCR tubes were left at room temperature for 10-20 minutes for the complementary resected ends of the insert and vector to anneal. Reaction mixtures were then used to transform 50 µl of chemicompetent T7Express *E. coli* cells by heat shock and transformants were plated on LB agar plates supplemented with ampicillin (100 µg/ml). Individual colonies were transferred to a replica plate and presence of insert was determined by colony PCR using the forward primer used for the assembly PCR and T7 terminator primer (GCTAGTTATTGCTCAGCGG). For expression, clones carrying an insert were used to inoculate 10 ml 2xYT bacterial cultures supplemented with ampicillin (100 µg/ml). The cultures were grown in a 50 ml centrifuge tube at 37 °C overnight with the tube lid slightly unscrewed and fixed with tape to ensure aeriation. Next day, protein expression was induced by the addition of IPTG (400 μM) for three hours at 37°C, after which cells were harvested by centrifugation. Cell pellets were then either frozen for future use or used directly for purification of peptides. Correct inserts were confirmed later by Sanger sequencing with the T7 terminator primer.

Cells were resuspended in 1 ml of lysis buffer: PBS, 20 mM imidazole, 1 mM TCEP, 1 mM EDTA, 0.1% Triton X-100, 0.2 mg/ml lysozyme, 1 mM AEBSF, 10 µg/ml DNase I. Lysate was incubated for 10 min at room temperature on a rotating mixer. Lysates were spun down in a 2 ml tube on a bench-top centrifuge at 15 000 g for 10 min and supernatant collected by aspiration. 200 µl of 50% (v/v) slurry of Ni-NTA agarose resin (Cube Biotech, !31103) was washed twice with 1 ml of MilliQ water and resuspended in 200 µl PBS. The resin was mixed with the soluble lysate and incubated on a rotating mixer for 5 min at room temperature, after which it was applied in two portions to a 0.5 ml micro-spin chromatography column and centrifugated for 1 min at 1000 x*g* to remove flow-through. Same centrifuge settings were also used for subsequent wash and elution steps. The resin was washed with a total of 1 ml of PBS + 20 mM imidazole containing 1 mM TCEP, followed by a 0.5 ml wash using the same buffer without any reducing agent. The second wash step is essential for the removal of any residual TCEP that can form undesired side-products upon reaction with the divinyl-heteroaryl linker. The GB1-BRC repeat was eluted with 0.5 ml PBS + 200 mM imidazole, and the elution immediately used for subsequent stapling reactions.

The eluted sample was split into two 250 µl parts. 2 mM DVT or DVP linker solution in DMSO was gradually titrated into the stapling reaction to achieve pseudo-dilution conditions. Different linker titration schemes were initially trialled and are shown in **Figure S2**. Most optimal linker titration was observed for reaction **h** (**Figure S2**) and was used for subsequent preps. At the same time, an identical volume of DMSO control without any linker was added to the other 250 µl peptide solution. 1 mM TCEP was added to the control reaction but not the stapling reaction to maintain free sulfhydryl groups in the control peptides. At the end of the titration, reactions were quenched with 2 mM DTT.

**Preparation of stapled peptides in un-tagged form**

Peptides were cloned in an identical fashion to the GB1-peptide-$His_8$ constructs, except different expression vectors, pOP3BT and pEXP-GB1, were used (Addgene #112603 and #112565, respectively). The vectors contain an N-terminal instead of a C-terminal His-tag. T7Express *E. coli* cells carrying the plasmids expressing GB1-fused BRC repeat were plated directly from glycerol stocks of sequence-verified clones onto LB agar supplemented with ampicillin (100 µg/ml) and grown overnight at 37 °C. Next day, cells were scraped to inoculate separate flasks containing 1 L of 2x YT medium supplemented with 100 µg/mL ampicillin. Cultures were grown at 37 °C until $OD_{600}$ of ~1, after which expression was induced with 400 µM IPTG for 3 h. Cells were resuspended in 25 mL of IMAC buffer A (50 mM Tris-HCl pH=8.0, 150 mM NaCl, 20 mM imidazole) and frozen. Later, cells were thawed and supplemented with DNase I (100 µL, 2 mg/mL) and AEBSF (1 mM), and lysed on an Emulsiflex C5 homogenizer (Avestin) or by sonication. Cell lysate was centrifuged at 40000 x*g* for 30 min and supernatant collected. GB1-BRC lysate was loaded on a 3 mL Ni-NTA agarose matrix (Cube Biotech), after which column matrix was washed with 10 column volumes Nickel Buffer A. GB1-BRC repeat was eluted with 12 ml nickel buffer B (50 mM Tris-HCl pH 8.0, 150 mM NaCl, 200 mM imidazole). The eluent was buffer exchanged back into nickel buffer A on a PD-10 desalting column (Cytiva). Buffer exchanged GB1-BRC fusion (~18 ml) was incubated with 100 µL of 2 mg/ml TEV protease overnight at 4°C. The GB1 tag was then removed from the solution by a second Ni-NTA affinity step, collecting the flow-through that contains the BRC peptide. The flow-through was acidified with HCl to pH 2-4 and acetonitrile was added to 10%, after which the solution was centrifuged at 10000 x*g* for 15 min and supernatant collected. The acidified flow-through was then applied to an ACE C8 300 4.6 x 250 mm semi-prep RP-HPLC column equilibrated with 10 % MeCN + 0.1% TFA and peptides were eluted with a 20 column volume gradient to 90 % MeCN + 0.1% TFA. BRC repeat peptides typically elute at 20-40% of the gradient. Peak fractions corresponding to the cleaved peptide were pooled and diluted 5x in PBS + 10 mM EDTA in a 50 ml centrifuge tube. A small stirrer bar was added to the tube, which was then placed on a magnetic stirrer. A syringe was filled with 20 mM linker in DMSO which was then gradually added to the mixture by piercing the centrifuge tube lid. To maintain pseudo-dilution conditions, linker was injected in 50 µl increments every two minutes, to a final concentration of 2 mM, ensuring at least 2x stoichiometric excess of linker over peptide. The reaction mixture was then quenched with 5 mM TCEP, filtered through a 0.45 µm filter and acidified with HCl to pH ~3. Stapled peptide was then purified by a on an ACE C18 300 4.6 x 250 mm semi-prep RP-HPLC column with a 0-100% gradient of A: 10 % MeCN + 0.1% TFA, B: 90 % MeCN + 0.1% TFA. Peak fractions containing the desired product were pooled and dried under vacuum.

**Purification of HumRadA22**

HumRadA22 is an archaeal RadA mutant with surface residues exchanged for the human Rad51 sequence and can be used as a bona fide mimic of monomeric human Rad51. The protein was prepared as described previously.[2]

**Purification of full-length human RAD51**

Full-length HsRAD51 was prepared based on a protocol developed at the lab of Prof Luca Pellegrini (Department of Biochemistry, University of Cambridge). *E. coli* BL21(DE3) Rosetta2 cells (Novagen) carrying a pRSF-Duet plasmid co-expressing wild-type *Hs*RAD51 and a BRC4 sequence fused to an N-terminal His-MBP tag were kindly provided by Dr Joseph

Maman. Cells were plated from a glycerol stock on LB agar supplemented with kanamycin (25 μg/mL) and chloramphenicol (34 μg/mL), and grown overnight at 37°C. Next day, cells were scraped and used to inoculate 1 L of 2xYT medium supplemented with same antibiotics. Cells were grown at 37°C with shaking at 200 RPM until an OD600 = 0.6, afer which they were cooled down to 18°C and expression induced with IPTG (400 μM) overnight. Cells were resuspended in 25 mL of buffer Ni-A-300 (50 mM Tris-HCl, pH 8.0, 300 mM NaCl, 20 mM imidazole, 1 mM TCEP) and frozen. Later, cells were thawed and supplemented with DNase I (100 μL, 2 mg/mL) and AEBSF (1 mM), and lysed on an Emulsiflex C5 homogenizer (Avestin) or by sonication. Cell lysate was spun down at 40 000 g for 30 min, after which the soluble fraction was loaded on a HisTrap HP 5 ml column (Cytiva). The column was washed with 8 CV Ni-A300 buffer, after which MBP-BRC4:RAD51 complex was co-eluted with buffer Ni-B-300 (50 mM Tris-HCl, pH 8.0, 300 mM NaCl, 200 mM imidazole, 1 mM TCEP). The sample was then diluted with Heparin-A buffer (20 mM HEPES pH 7.4, 50 mM NaCl, 1 mM EDTA, 1 mM TCEP) and loaded on a HiTrap Heparin HP 5 ml column. During this step, RAD51 oligomerises on the heparin matrix, which acts as a DNA mimic, and dissociates from the MBP-BRC4 fusion, which is removed in flow-through and wash steps. Column was washed with 8 CV Heparin-A buffer, after which the protein was eluted with a 20 CV, 0-100% linear gradient of Heparin-B buffer (20 mM HEPES pH 7.4, 1 M NaCl, 1 mM EDTA, 1 mM TCEP). RAD51 was concentrated, flash-frozen with liquid nitrogen and stored for future use.

**Fluorescence polarisation competition assay**

FP competition assay was a modified version of a protocol described previously.[2] Fluorescein-labelled *Hs*BRC4 probe for this assay was kindly provided by Dr Laurens Lindenburg (Hollfelder group, Department of Biochemistry, University of Cambridge). Black 384-well flat-bottom microplates (Corning, 3821) were used with a 40 μl final reaction volume in all measurements. Following buffer conditions were used: 20 mM CHES pH 9.5, 150 mM NaCl, 0.1% BSA, 0.1% Tween-20. Each reaction contained 100 nM HumRadA22 and 10 nM BRC4-fluorescein. Two-fold serial dilutions of stapled peptides were added to the reactions. A free probe control reaction containing only 10 nM BRC4-fluorescein was used to calibrate gain and focal height. FP measurements were performed on a Pherastar FX (BMG Labtech) plate reader equipped with an FP 485-520-520 optic module. Binding curves were fitted using the four-parameter logistic model with a variable Hill slope using Prism software (Graphpad). Regression fitting was performed using the least squares optimisation algorithm. $K_D$ values were estimated from the fitted $IC_{50}$ parameters using a previously reported equation.[6]

**Isothermal titration calorimetry**

Peptides were resuspended in MilliQ water to 10 times the desired final concentration. This was then diluted 10x with the ITC buffer to obtain the final titrant solution (20 mM CHES pH 9.5, 150 mM NaCl, 0.1% Tween-20). HumRadA22 was buffer-exchanged on a NAP-5 desalting column (Cytiva) into ITC buffer and protein concentration was adjusted to 10:9 of the desired final value. One ninth volume of MilliQ water was added to the solution to bring the protein concentration to the desired final value, while maintaining identical buffer:MilliQ volume proportions in both the syringe and the cell. ITC was carried out using a Microcal ITC200 instrument at 25°C with a 5.00 μCal reference power DP value, stirring speed of 500-750 rpm, 2 sec filter period. Injection spacing, speed and volume, cell/syringe concentrations as well as the number of injections were adjusted for each peptide and its binding properties. ITC data were fitted using a single-site binding model using the Microcal ITC data analysis program in the Origin 7.0 package. Data points affected by baseline spikes were omitted from the analysis.

**Circular dichroism spectroscopy**

Dried peptides were dissolved in MilliQ water to 0.3 mg/ml, and then two-fold diluted in 20 mM sodium phosphate, pH 7.4, giving a final solution of 0.15 mg/ml peptide in 10 mM sodium phosphate. CD spectra of selected peptides were recorded on an AVIV 410 circular dichroism spectropolarimeter using a 1 mm path length quartz cuvette. Measurements were done at 25 °C, with a 185-260 nm range, 1 nm bandwidth, 5 s averaging time and 0.3 s settling time. Spectra were prepared as smoothed average of three scans and normalised against blank solvent.

**Electrophoretic mobility shift assay (EMSA)**

The ability of linear and stapled BRC repeat peptides to dissociate RAD51-ssDNA nucleofilament was evaluated using an electrophoretic mobility shift assay (EMSA). RAD51 DNA-binding reactions (40 μl) were set up in 50 mM HEPES pH 7.4, 150 mM NaCl, 10 mM MgAc2, 2 mM CaCl2, 1 mM TCEP, 1 mM ATP. 5 μM full-length human RAD51 was incubated with varying concentrations of BRC repeats for 10 min at room temperature, followed by the addition of 100 nM fluorescently labelled FAM-dT60 oligonucleotide, and further incubation at 37 °C for 10 min. Control reactions were set up with free FAM-dT60 probe and FAM-dT60 Pantelejevs, T. Materials and methods 120 + 5 μM RAD51. 10 μl of reactions were then loaded on a 1xTBE non-denaturing acrylamide gel (5%) and run at 100 V for 1:30 h at 4 °C. The gel was directly visualized on a Typhoon FLA 9000 imager (GE Healthcare) using FAM channels.

**X-ray crystallography**

Stapled peptide complexes were re-constituted from purified peptides and HumRadA22. Peptides were added at a 1.5 stoichiometric excess to HumRadA22 in its size-exclusion buffer, to a final concentration of 0.75 and 0.5 mM for the peptide and protein, respectively. ADP and $MgCl_2$ were added to the protein solutions in for some of the complexes (see **Table S1**). Crystallisation screening was done in 96-well MRC plates using the sitting-drop vapor diffusion technique and a variety of commercial crystallisation screens. A Mosquito liquid handling robot (TTP Labtech) was used to dispense protein and reservoir solutions in sub-microlitre volumes. Typically the two sitting drops contained 200 or 400 nl of protein solution and 200 nl of crystallisation solution, while the reservoir contained 80 μl of crystallisation solution. Plates were stored at 17 °C in a RockImager crystallisation hotel (Formulatrix) and imaged regularly. Crystal hits were flash-frozen in liquid nitrogen using cryoloops. Additional cryoprotectant was not added before freezing of crystal hits. Diffraction data were collected on Diamond Light Source (Harwell, UK) MX beamlines. Full native datasets with goniometer sweeps of at least 180° were collected to ensure completeness of diffraction data. Molecular replacement phasing method was used with the apo HumRadA22 structure (PDB: 5KDD) as a search model. Molecular replacement was done with Phaser.[7] The structures were refined without BRC repeats first and the peptides were built into the clearly visible electron density manually. Manual refinement was done in Coot[8] and automated refinement with phenix.refine[7] and autoBUSTER.[9] Crystallisation conditions, as well as data collection and refinement statistics, are provided in **Table S2**. The coordinates have been deposited in the Protein Data Bank under accession codes 8C3J (**SP2**), 8BR9 (**SP24**) and 8C3N (**SP30**).

**Cell line**

U2OS cell line (ATCC, HTB-96) was used in this study. They were cultured in DMEM medium (Sigma-Aldrich, D6429) containing 10% fetal bovine serum (Sigma-Aldrich, F7524) and 100 U/mL penicillin/streptomycin (Sigma-Aldrich, P4333) at 37 °C and 5% CO2

**Immunofluorescence**

50000 cells per well were seeded in twelve-well plates containing one round coverslips in each well. After 2-3 days, medium was replaced by fresh medium containing or not **SP31** (40 μM). Cells were incubated for an hour before they were irradiated (3 Gy) and allowed to recover for 3 hours before being fixed. Non-irradiated cells treated in a similar way were used as a control.

RAD51 foci were detected following a protocol previously described.[1] In summary, after being washed with PBS, cells were fixed with paraformaldehyde, 4% in PBS (freshly made from paraformaldehyde 32% Aqueous sol. EM GRADE, Electron Microscopy Science, 15714-S) for 15 minutes at room temperature. After washing with PBS, cells were permeabilisated with Triton X-100 (Sigma-Aldrich, T8787), 0.5% in PBS for 7 min at room temperature. Cells were washed again and blocked with 1% BSA (Sigma-Aldrich, A9647) in PBS for at least an hour at room temperature. RAD51 antibody (Santa Cruz Biotechnology, sc-8349) was diluted 1:100 in blocking solution and added to the fixed cells for around 2 hours at room temperature. After washing with PBS, cells were incubated with anti-rabbit IgG Alexa Fluor 488 (Life Technologies, A21206) diluted 1:500 in the blocking solution for around an hour at room temperature. Finally, cells were washed with PBS and slides were prepared by adding a drop of mounting medium with DAPI (ProLong Gold Antifade Mountant with DAPI, ThermoFisher, P36935) and stored at 4 °C. Images were taken using a Nikon Eclipse e-400 microscope with a 40x objective and the software SlideBook 6. RAD51 foci were counted using CellProfiler 4.0.6 and GraphPad Prism 9.4.1.

### pPEPT1 vector

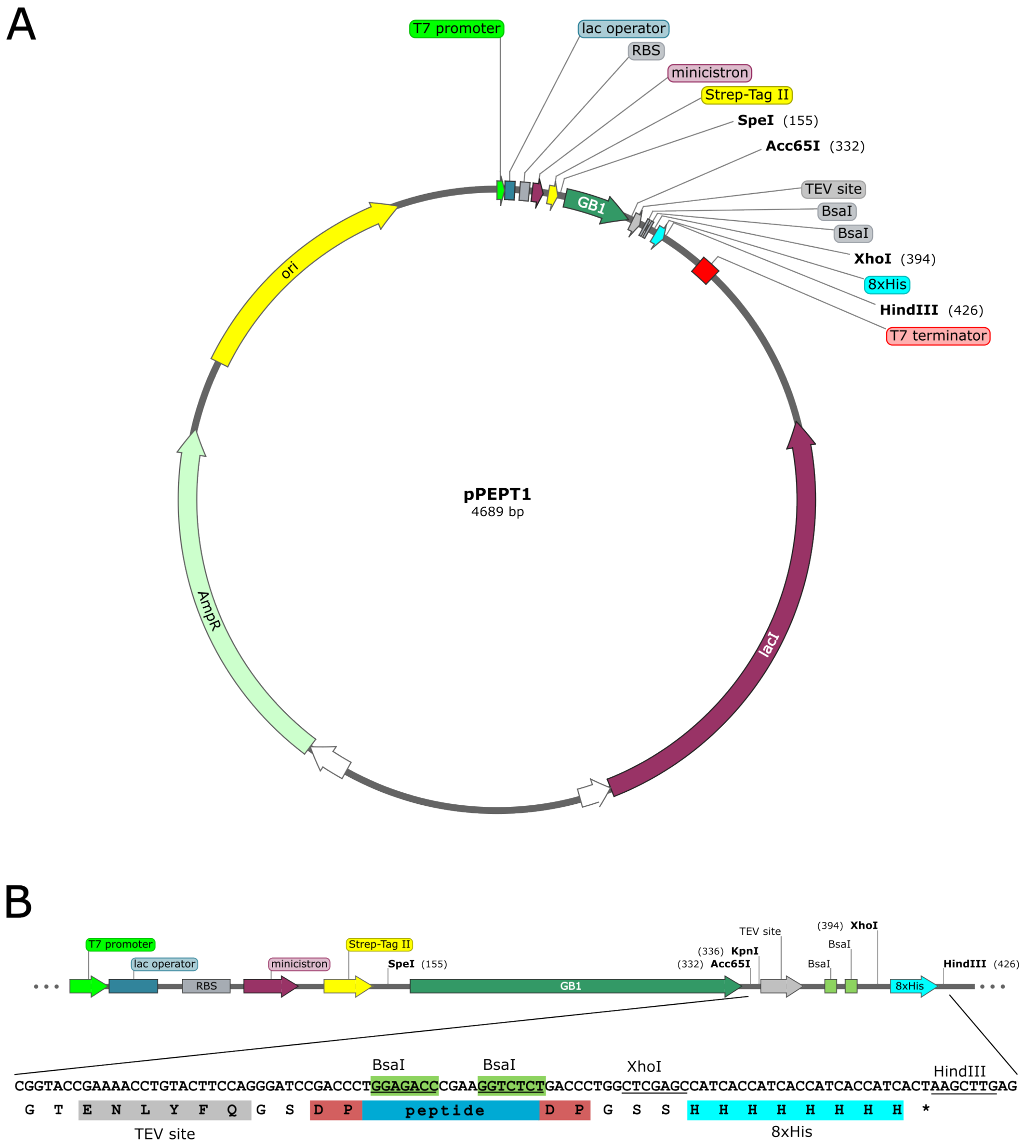

**Figure S1.** Map of the pPEPT1 plasmid. (A) circular plasmid map of the pPEPT1 vector with key features and unique restrictions sites in the fusion part and in the multiple cloning site. (B) Focused view of the DNA and protein features of the part where the peptide-encoding sequences are inserted.

### Optimisation of the small-scale stapling reaction

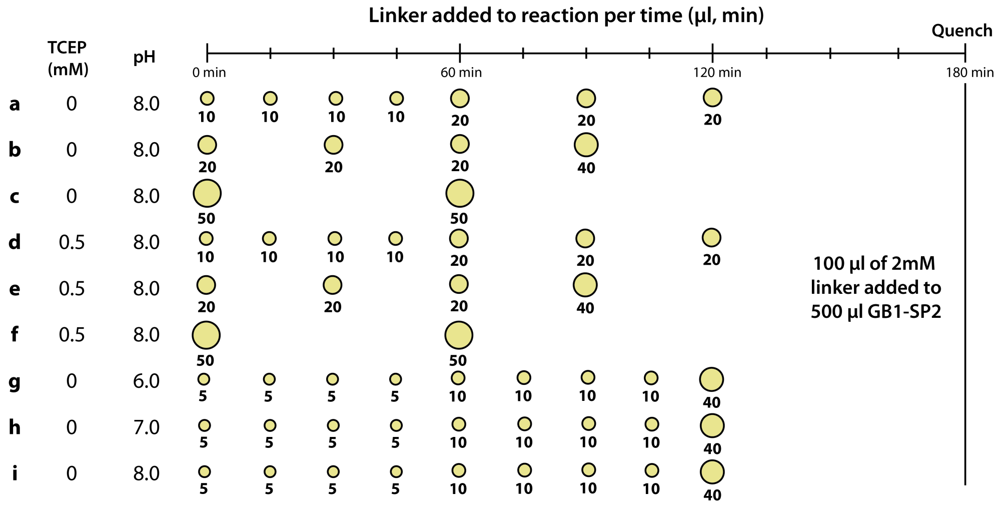

**Figure S2.** Optimisation of small-scale stapling reaction. Lower-case letters a-i represent different conditions. For each condition, the volumes (µl) of linker addition are depicted at appropriate time-points under the yellow circles, are which is relative to the volume for more visual interpretation of the experiments. pH and presence of TCEP are also indicated. ESI-MS mass spectra for each condition are shown below.

### X-ray crystallography statistics

| Complex | SP2:HumRadA22 | SP24:HumRadA22 | SP30:HumRadA22 |
|---|---|---|---|
| **Protein** | 0.5 mM SP2:HumRadA22 in 20 mM CHES pH 9.5, 100 mM NaCl | 0.5 mM SP24:HumRadA22 in 20 mM CHES pH 9.5, 100 mM NaCl, 20 mM ADP/$MgCl_2$ | 0.5 mM SP30:HumRadA22 in 20 mM CHES pH 9.5, 100 mM NaCl, 20 mM ADP/$MgCl_2$ |
| **Condition** | 0.1 M $Na_3$Citrate pH 4.2, 20% w/v PEG 1K, 0.2 M $Li_2SO_4$ | 14% w/v PEG 4000, 6% v/v MPD, 0.1M Na K Phos pH 6.2 | 14% w/v PEG 4000, 6% v/v MPD, 0.1M Na K Phos pH 6.2 |
| **Protein:well solution (nl:nl)** | 200:200 | 400:200 | 200:200 |
| **PDB code** | 8C3J | 8BR9 | 8C3N |
| **Data collection and processing** | | | |
| Beamline | DLS i03 | DLS i04 | DLS i04 |
| Wavelength (Å) | 0.9762 | 0.9795 | 0.9795 |
| Space group | P 41 21 2 | P 21 21 2 | P 21 21 2 |
| a, b, c (Å) | 112.56 112.56 140.79 | 140.46 38.64 43.53 | 143.13 38.01 43.92 |
| α, β, γ (°) | 90.00 90.00 90.00 | 90.00 90.00 90.00 | 90.00 90.00 90.00 |
| Resolution range (Å) | 87.92 - 3.02 (3.02 - 3.07) | 70.23 - 1.63 (1.66 - 1.63) | 1.25 - 41.99 (1.25 - 1.27) |
| $R_{meas}$ | 0.242 (3.751) | 0.059 (4.504) | 0.046 (1.597) |
| Completeness (%) | 100.0 (100.0) | 98.7 (99.2) | 99.6 (95.3) |
| Reflections | 170095 / 13512 | 239726 / 30303 | 513195 / 67481 |
| Redundancy | 12.8 (13.8) | 7.9 (7.8) | 7.6 (4.6) |
| $\langle I/\sigma(I) \rangle$ | 10.0 (0.9) | 12.0 (0.3) | 15.8 (0.8) |
| CC½ | 1.0 (0.3) | 1.0 (0.3) | 1.0 (0.4) |
| **Refinemnt** | | | |
| $R_{cryst}/R_{free}$ | 0.228 / 0.254 | 0.260 / 0.270 | 0.196 / 0.210 |
| Resolution range (Å) | 69.29 - 3.02 | 70.23 - 1.62 | 36.74 - 1.24 |
| Reflections in work / test set | 17515 / 897 | 27946 / 1450 | 64107 / 3361 |
| Number of atoms | 3834 | 2041 | 2373 |
| Mean / Wilson B-factor (Å$^2$) | 95.6 / 79.5 | 63.3 / 34.4 | 27.9 / 17.8 |
| Ramachandran favoured/allowed/outliers (%) | 97.22 / 2.57 / 0.21 | 99.17 / 0.83 / 0.00 | 99.15 / 0.85 / 0.00 |
| RMSD bonds (Å) | 0.017 | 0.013 | 0.014 |
| RMSD angles (°) | 1.66 | 1.69 | 1.61 |

**Table S1**. Crystallographic data collection and refinement. Values in parentheses are for the high-resolution cell.

### Rad51 foci inhibition data

**A**

#### Experiment 1

| Control noIR | | |
|---|---|---|
| Foci = 0 | 552 | 54.2 |
| Foci = 1 | 90 | 8.8 |
| Foci = 2 | 47 | 4.6 |
| Foci = 3 | 40 | 3.9 |
| Foci = 4 | 44 | 4.3 |
| Foci 5-9 | 149 | 14.6 |
| Foci > 10 | 96 | 9.4 |
| Number of cells | 1018 | 100.0 |

| Control 3Gy 3h | | |
|---|---|---|
| Foci = 0 | 458 | 48.3 |
| Foci = 1 | 90 | 9.5 |
| Foci = 2 | 59 | 6.2 |
| Foci = 3 | 45 | 4.7 |
| Foci = 4 | 34 | 3.6 |
| Foci 5-9 | 116 | 12.2 |
| Foci > 10 | 146 | 15.4 |
| Number of cells | 948 | 100.0 |

| Stpl31 noIR | | |
|---|---|---|
| Foci = 0 | 1015 | 83.3 |
| Foci = 1 | 91 | 7.5 |
| Foci = 2 | 44 | 3.6 |
| Foci = 3 | 30 | 2.5 |
| Foci = 4 | 8 | 0.7 |
| Foci 5-9 | 28 | 2.3 |
| Foci > 10 | 2 | 0.2 |
| Number of cells | 1218 | 100.0 |

| Stpl31 3Gy 3h | | |
|---|---|---|
| Foci = 0 | 1029 | 88.9 |
| Foci = 1 | 47 | 4.1 |
| Foci = 2 | 32 | 2.8 |
| Foci = 3 | 14 | 1.2 |
| Foci = 4 | 4 | 0.3 |
| Foci 5-9 | 24 | 2.1 |
| Foci > 10 | 8 | 0.7 |
| Number of cells | 1158 | 100.0 |

#### Experiment 2

| Control noIR | | |
|---|---|---|
| Foci = 0 | 1279 | 70.3 |
| Foci = 1 | 156 | 8.6 |
| Foci = 2 | 103 | 5.7 |
| Foci = 3 | 86 | 4.7 |
| Foci = 4 | 47 | 2.6 |
| Foci 5-9 | 122 | 6.7 |
| Foci > 10 | 26 | 1.4 |
| Number of cells | 1819 | 100.0 |

| Control 3Gy 3h | | |
|---|---|---|
| Foci = 0 | 802 | 57.2 |
| Foci = 1 | 92 | 6.6 |
| Foci = 2 | 63 | 4.5 |
| Foci = 3 | 69 | 4.9 |
| Foci = 4 | 53 | 3.8 |
| Foci 5-9 | 183 | 13.0 |
| Foci > 10 | 141 | 10.0 |
| Number of cells | 1403 | 100.0 |

| Stpl31 noIR | | |
|---|---|---|
| Foci = 0 | 1103 | 72.4 |
| Foci = 1 | 174 | 11.4 |
| Foci = 2 | 97 | 6.4 |
| Foci = 3 | 56 | 3.7 |
| Foci = 4 | 35 | 2.3 |
| Foci 5-9 | 54 | 3.5 |
| Foci > 10 | 4 | 0.3 |
| Number of cells | 1523 | 100.0 |

| Stpl31 3Gy 3h | | |
|---|---|---|
| Foci = 0 | 821 | 63.6 |
| Foci = 1 | 108 | 8.4 |
| Foci = 2 | 54 | 4.2 |
| Foci = 3 | 54 | 4.2 |
| Foci = 4 | 46 | 3.6 |
| Foci 5-9 | 129 | 10.0 |
| Foci > 10 | 79 | 6.1 |
| Number of cells | 1291 | 100.0 |

#### Experiment 3

| Control noIR | | |
|---|---|---|
| Foci = 0 | 754 | 59.7 |
| Foci = 1 | 120 | 9.5 |
| Foci = 2 | 71 | 5.6 |
| Foci = 3 | 55 | 4.4 |
| Foci = 4 | 51 | 4.0 |
| Foci 5-9 | 138 | 10.9 |
| Foci > 10 | 75 | 5.9 |
| Number of cells | 1264 | 100.0 |

| Control 3Gy 3h | | |
|---|---|---|
| Foci = 0 | 581 | 51.2 |
| Foci = 1 | 93 | 8.2 |
| Foci = 2 | 47 | 4.1 |
| Foci = 3 | 30 | 2.6 |
| Foci = 4 | 40 | 3.5 |
| Foci 5-9 | 144 | 12.7 |
| Foci > 10 | 199 | 17.5 |
| Number of cells | 1134 | 100.0 |

| Stpl31 noIR | | |
|---|---|---|
| Foci = 0 | 825 | 74.3 |
| Foci = 1 | 114 | 10.3 |
| Foci = 2 | 61 | 5.5 |
| Foci = 3 | 39 | 3.5 |
| Foci = 4 | 17 | 1.5 |
| Foci 5-9 | 38 | 3.4 |
| Foci > 10 | 16 | 1.4 |
| Number of cells | 1110 | 100.0 |

| Stpl31 3Gy 3h | | |
|---|---|---|
| Foci = 0 | 1014 | 78.4 |
| Foci = 1 | 114 | 8.8 |
| Foci = 2 | 36 | 2.8 |
| Foci = 3 | 29 | 2.2 |
| Foci = 4 | 15 | 1.2 |
| Foci 5-9 | 59 | 4.6 |
| Foci > 10 | 27 | 2.1 |
| Number of cells | 1294 | 100.0 |

**B**

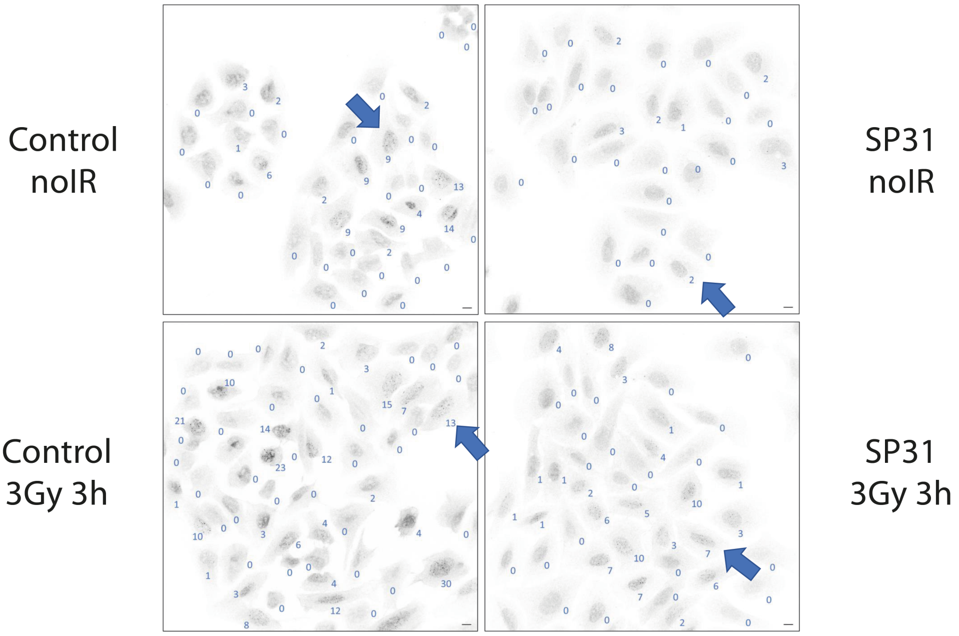

**Figure S3**. (A) Quantification of RAD51 foci in U2OS cells from three independent experiments. (B) Representative IF images depicting RAD51 signal. Representative cells shown in **Error! Reference source not found.**B are indicated with an arrow a nd they were chosen as the sixth highest RAD51 foci for each of the conditions. Foci counts determined automatically using CellProfiler software are provided next to each cell. Scale bar 10 mm.

### Oligonucleotides used for cloning of peptides

| Insert | Vector | Res. enzyme(s) | No | Sequence |
|---|---|---|---|---|
| SP1 | pPEPT1 | *BsaI* | 1 | TTCCAGGGATCCGACCCTTCTTGTGAAGCGCTGCAGAAGGCGTGTAAACTGTTCTC |
| | | | 2 | ATGGCTCGAGCCAGGGTCAGAGATGTTTTCGATGTCGGAGAACAGTTTACACGCCTT |
| SP2 | pOP3BT | *BamHI/XhoI* | 1 | CCTGTACTTCCAGGGATCCGTTAACTCTTCTGCGTTCTCCG |
| | | | 2 | CAGTTTTTTACCAGACGCGGTAGAGAAACCGGAGAACGCAGAAGAGTTAA |
| | | | 3 | CCGCGTCTGGTAAAAAACTGAACGTTTCTTGCGAAGCGCTGCAGAAAGCG |
| | | | 4 | CAAGCTTAGCTCGAGCCAGAGATGTTTTCGATGTCAGAGAACAGTTTGCACGCTTTCTGCAGCGCTT |
| | pPEPT1 | *BsaI* | 1 | TTCCAGGGATCCGACCCTGTTAACTCTTCTGCGTTCTCCGGTTTCTCTACCGCGTCTGGTAAAAAACTGAACGTTTCTTGCGA |
| | | | 2 | ATGGCTCGAGCCAGGGTCAGAGATGTTTTCGATGTCAGAGAACAGTTTGCACGCTTTCTGCAGCGCTTCGCAAGAAACGTTCAGTTTT |
| SP7 | pPEPT1 | *BsaI* | 1 | TTCCAGGGATCCGACCCTGTTAACTCTTCTGCGTTCTCTGGTTTCTGCACCGCGTCTGGTAAAAAACTGAACGTTTCTTGCGA |
| | | | 2 | ATGGCTCGAGCCAGGGTCAGAGATGTTTTCGATGTCGGAGAACAGTTTAACCGCCTTCTGCAGCGCTTCGCAAGAAACGTTCAGTTTT |
| SP8 | pPEPT1 | *BsaI* | 1 | TTCCAGGGATCCGACCCTGTTAACTCTTCTGCGTTCTCTGGTTTCTCTACCGCGTCTGGTAAAAAACTGAACGTTTCTACCTGTG |
| | | | 2 | ATGGCTCGAGCCAGGGTCAGAGATGTTTTCGATGTCGGAGAACAGACAAACCGCCTTCTGCAGCGCACAGGTAGAAACGTTCAGT |
| SP9 | pPEPT1 | *BsaI* | 1 | TTCCAGGGATCCGACCCTGTTAACTCTTCTGCGTTCTCTGGTTTCTCTACCGCGTCTGGTAAAAAACTGAACGTTTCTACCGAAG |
| | | | 2 | ATGGCTCGAGCCAGGGTCAGAGATGTTTTCGATGTCACAGAACAGTTTAACCGCCTTACACAGCGCTTCGGTAGAAACGTTCAGT |
| SP10 | pPEPT1 | *BsaI* | 1 | TTCCAGGGATCCGACCCTGCGTGTGGTTTCTCTACCGCGTCTGGTAAAAAACTGAACGTTTCTTGTGAAGCGCTGCAG |
| | | | 2 | ATGGCTCGAGCCAGGGTCAGAGATGTTTTCGATGTCGGAGAACAGTTTAACCGCCTTCTGCAGCGCTTCACAAGAAA |
| SP11 | pPEPT1 | *BsaI* | 1 | TTCCAGGGATCCGACCCTTCTTGTTTCTCTACCGCGTCTGGTAAAAAACTGAACGTTTCTTGTGAAGCGCTGCAGA |
| | | | 2 | ATGGCTCGAGCCAGGGTCAGAGATGTTTTCGATGTCGGAGAACAGTTTAACCGCCTTCTGCAGCGCTTCACAAGAA |
| SP12 | pPEPT1 | *BsaI* | 1 | TTCCAGGGATCCGACCCTTCTTCTGCGTGTTCTGGTTTCTCTACCGCGTCTGGTAAAAAACTGAACGTTTCTACCGAAGCGT |
| | | | 2 | ATGGCTCGAGCCAGGGTCAGAGATGTTTTCGATGTCGGAGAACAGTTTAACCGCCTTCTGACACGCTTCGGTAGAAACGTTC |
| SP13 | pPEPT1 | *BsaI* | 1 | TTCCAGGGATCCGACCCTGTTAACTGTTCTGCGTTCTCTGGTTTCTCTACCGCGTCTGGTAAAAAACTGAACGTTTCTACCGAAG |
| | | | 2 | ATGGCTCGAGCCAGGGTCAGAGATGTTTTCGATGTCGGAGAACAGTTTAACCGCCTTACACAGCGCTTCGGTAGAAACGTTCAGT |
| SP14 | pPEPT1 | *BsaI* | 1 | TTCCAGGGATCCGACCCTGTTAACTGTTCTGCGTTCTCTGGTTTCTCTACCGCGTCTG |
| | | | 2 | ATGGCTCGAGCCAGGGTCAGAAACACACAGTTTTTTACCAGACGCGGTAGAGAAACCA |
| SP16 | pPEPT1 | *BsaI* | 1 | TTCCAGGGATCCGACCCTGCGTGTTCTGGTTTCTCTACCGCGTCTGGTAAAAA |
| | | | 2 | ATGGCTCGAGCCAGGGTCAGAACAAACGTTCAGTTTTTTACCAGACGCGGTAGA |
| SP24 | pEXP-Nhis-GB1 | *BsaI/HindIII* | 1 | ACCTGTACTTCCAGGGATCCGGTTTCTCTACCGCGTCTGGT |
| | | | 2 | GCAGCGCCTGGCAAGAAACGTTCAGTTTTTTACCAGACGCGGTAGAGAAA |
| | | | 3 | CTTGCCAGGCGCTGCAGAAAGCGTGCAAACTGTTCTCTGGTAGCCATCAT |
| | | | 4 | TGGTGATGGTGATGGCTCGAGCCCCAATGATGGCTACCAGAGAACAG |
| SP30 | pEXP-Nhis-GB1 | *BsaI/HindIII* | 1 | TTCTAATACGACTCACTATAGGTACCGAAAACCTGTACTTCC |
| | | | 2 | GCCGCTCGCGGTGCTAAAGCCGCTGCAGCCCTGGAAGTACAGGTTTTCGGTACCTATAGTGAGTC |
| | | | 3 | CCGCGAGCGGCAAAAAACTGAACGTGAGCACCCAGGCGTGCCAGA |
| | | | 4 | CTATAGAATACTCAAGCTTAGCCGCTAAACAGTTTCACCGCTTTCTGGCACGCCTGGGT |

**Table S2**. Oligonucleotides used for the cloning of peptide constructs.

### Mass spectra

#### Mass spectra of reaction optimisation conditions with SP2

| Species | mass (calc), Da |
|---|---|
| GB1-**SP2**-DVT (-Met1) | 14773.97 |
| GB1-**SP2**-DVT (-Met1, -Ser2) | 14686.89 |
| GB1-**SP2**-$(DVT)_2$ (-Met1) | 14994.2 |
| GB1-**SP2**-$(DVT)_2$-TCEP (-Met1) | 15244.38 |
| GB1-**SP2**-$(DVT)_2$-$(TCEP)_2$ (-Met1) | 15494.57 |

##### Reaction **a**

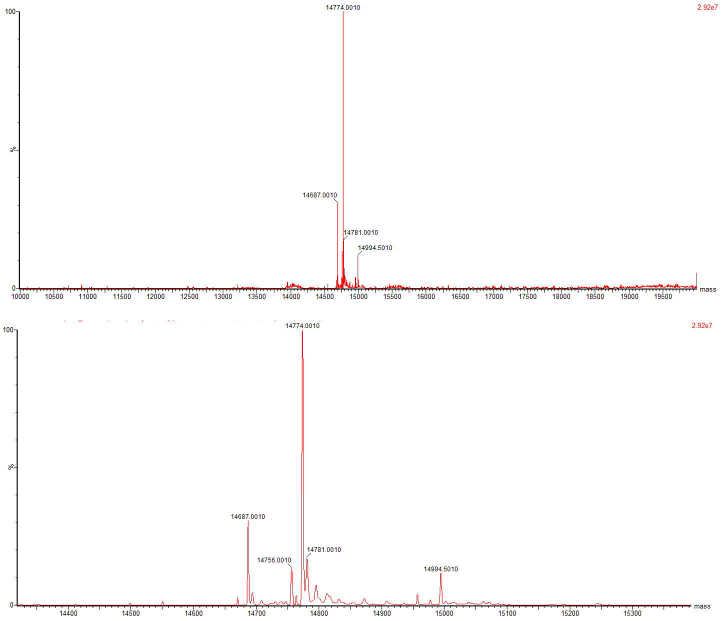

**Figure S4**. MS analysis of the reaction products of reaction **a**, for which the conditions are described in **Figure S2**

##### Reaction **b**

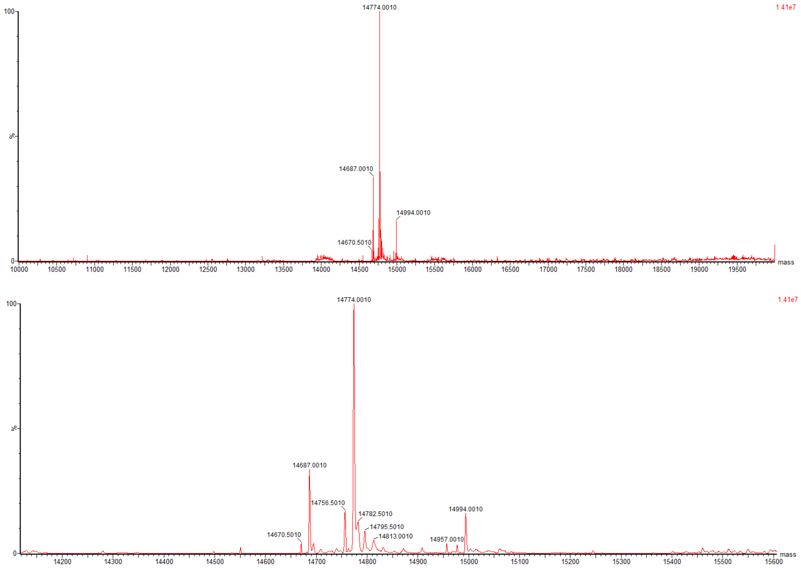

**Figure S5**. MS analysis of the reaction products of reaction **d**, for which the conditions are described in **Figure S2**

Reaction **c**

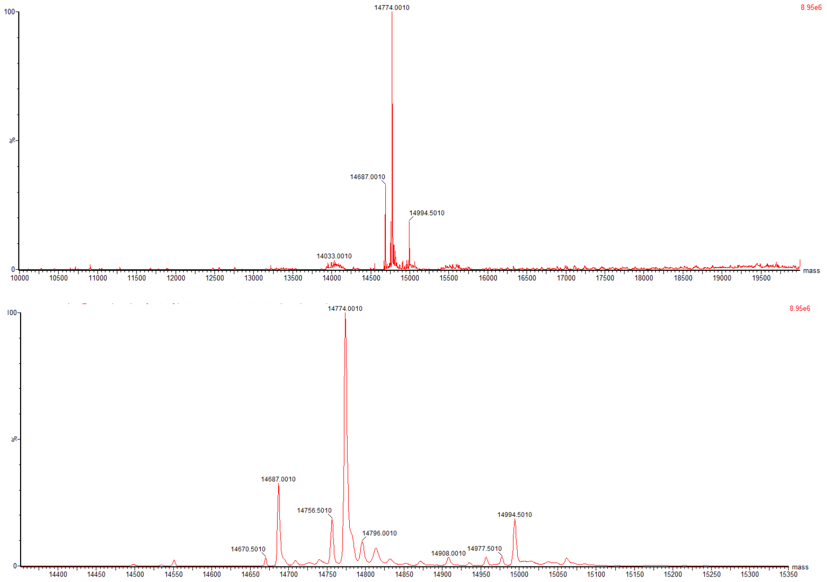

**Figure S6**. MS analysis of the reaction products of reaction **c**, for which the conditions are described in **Figure S2**

Reaction **d**

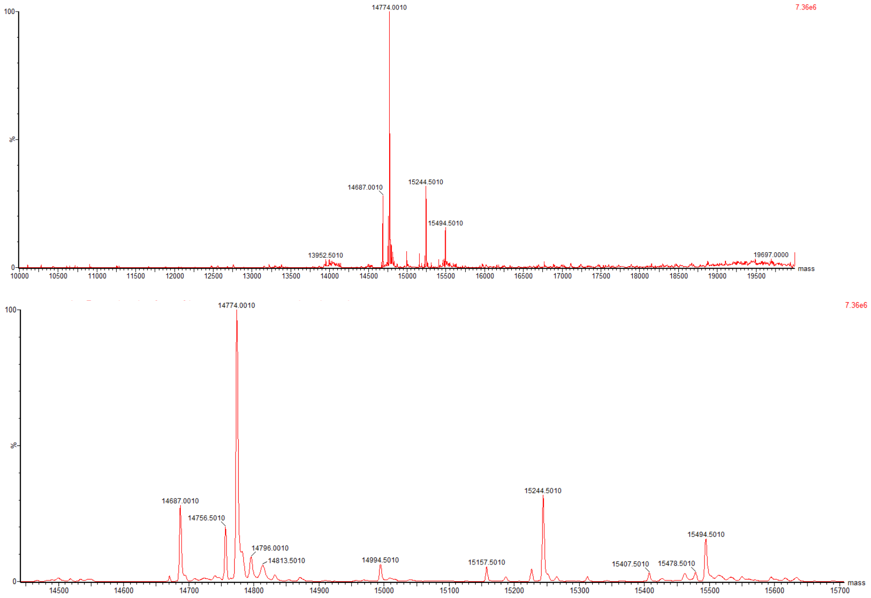

**Figure S7**. MS analysis of the reaction products of reaction **d**, for which the conditions are described in **Figure S2**

#### Reaction **e**

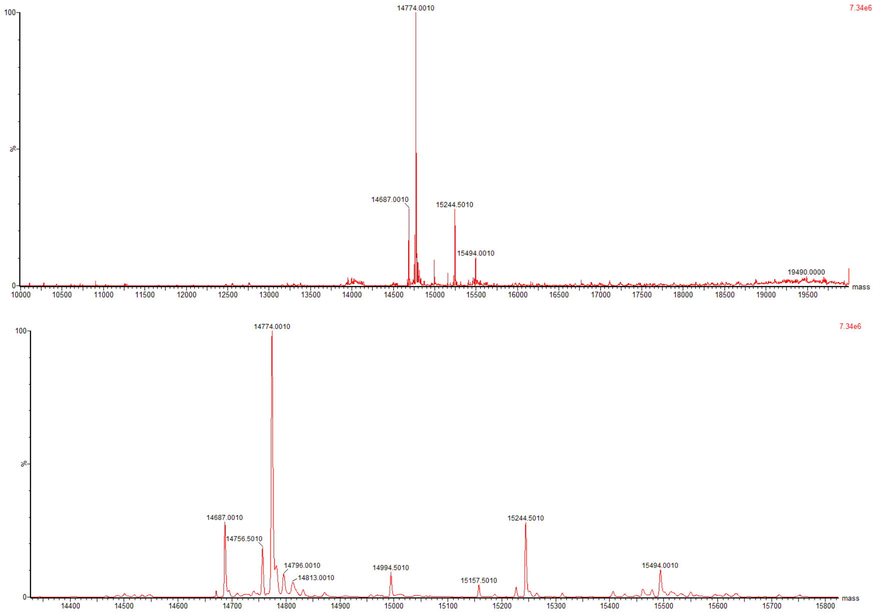

**Figure S8**. MS analysis of the reaction products of reaction **e**, for which the conditions are described in **Figure S2**

#### Reaction **f**

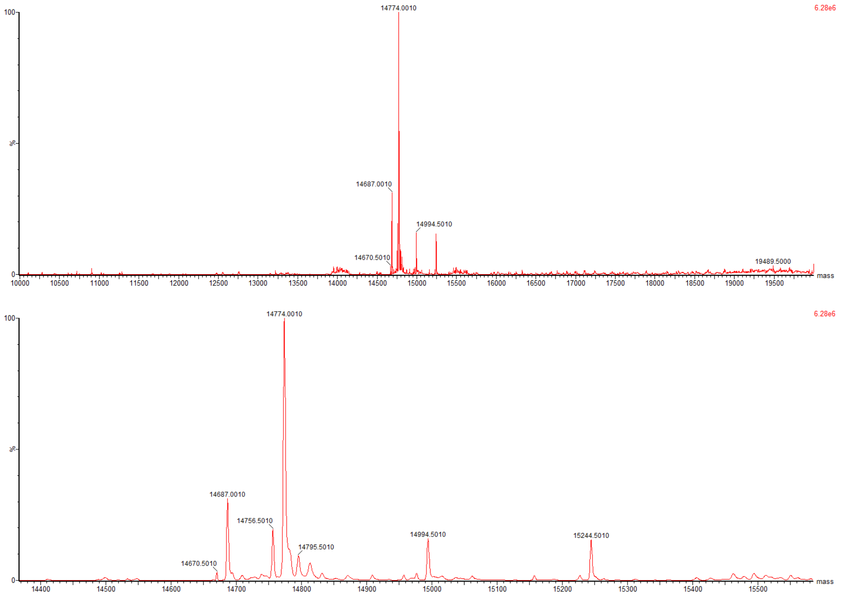

**Figure S9**. MS analysis of the reaction products of reaction **f**, for which the conditions are described in **Figure S2**

#### Reaction **g**

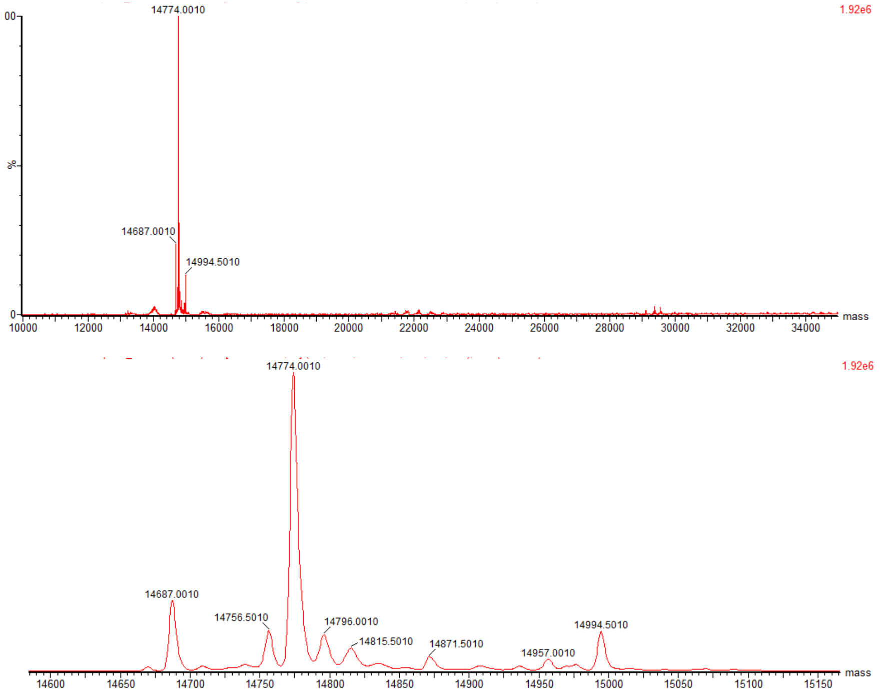

**Figure S10**. MS analysis of the reaction products of reaction **g**, for which the conditions are described in **Figure S2**

#### Reaction **h**

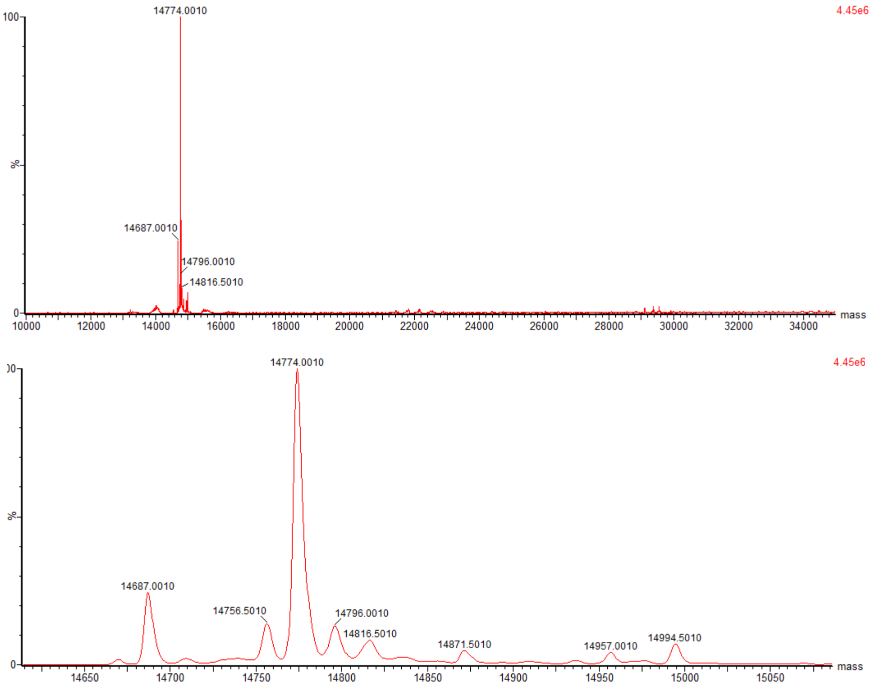

**Figure S11**. MS analysis of the reaction products of reaction **h**, for which the conditions are described in **Figure S2**

### Mass spectra of representative GB1-fused stapled peptides

## SP10

| Species | mass (calc), Da | mass (obs), Da |
|---|---|---|
| GB1-**SP10**-DVT (-Met1) | 14251.46 | 14251.5 |
| GB1-**SP10**-DVT (-Met1, -Ser2) | 14164.38 | 14164.5 |
| GB1-**SP10**-(DVT)$_2$ (-Met1) | 14471.69 | 14471.5 |

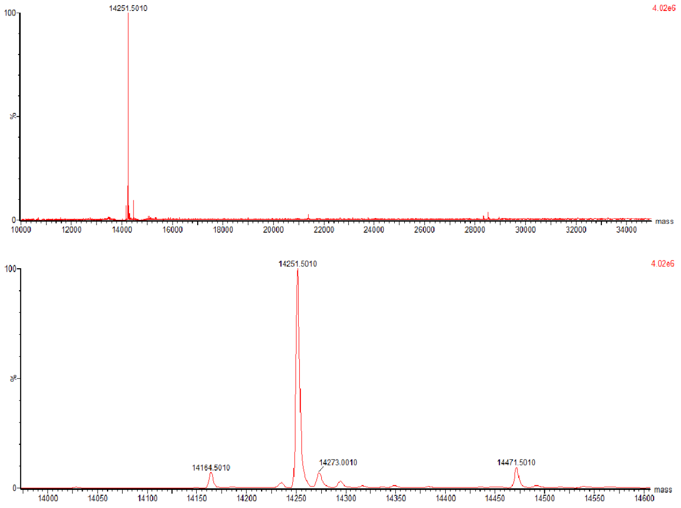

**Figure S12**. MS analysis of the GB1-fused peptide **SP10**.

## SP11

| Species | mass (calc), Da | mass (obs), Da |
|---|---|---|
| GB1-**SP11**-DVT (-Met1) | 14210.4 | 14210.2 |
| GB1-**SP11**-DVT (-Met1, -Ser2) | 14123.33 | 14122.9 |
| GB1-**SP11**-(DVT)$_2$ (-Met1) | 14430.63 | 14430.4 |

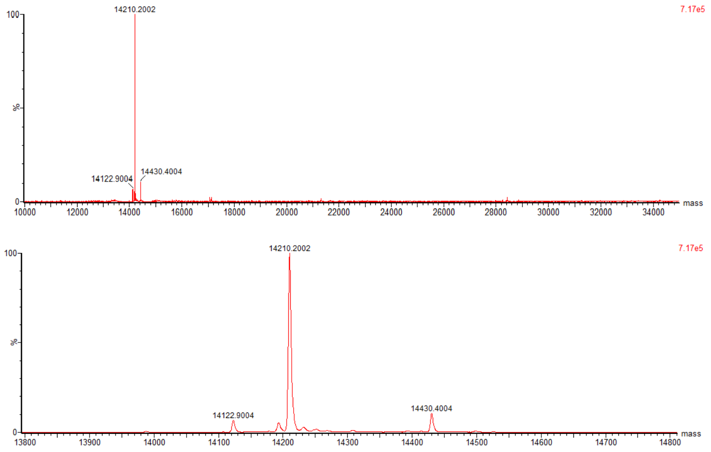

**Figure S13**. MS analysis of the GB1-fused peptide **SP11**.

## SP12

| Species | mass (calc), Da | mass (obs), Da |
|---|---|---|
| GB1-**SP12**-DVT (-Met1) | 14500.64 | 14500.5 |
| GB1-**SP12**-DVT (-Met1, -Ser2) | 14720.87 | 14720.5 |
| GB1-**SP12**-$(DVT)_2$ (-Met1) | 14413.56 | 14413.5 |

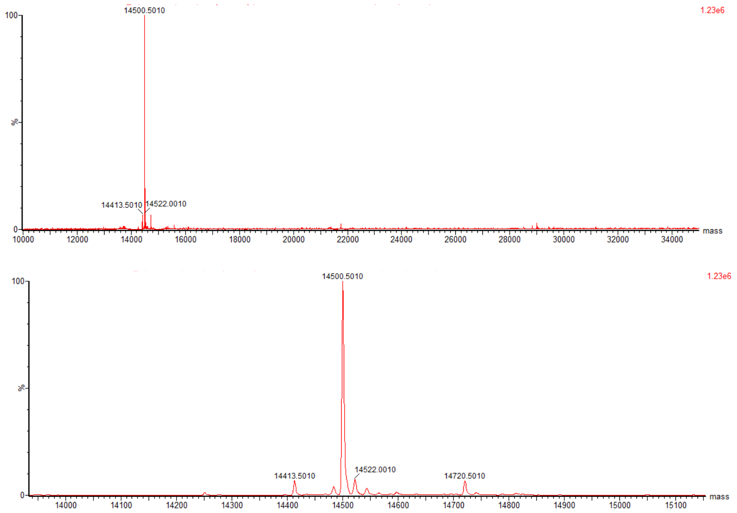

**Figure S14**. MS analysis of the GB1-fused peptide **SP12.**

## SP15

| Species | mass (calc), Da | mass (obs), Da |
|---|---|---|
| GB1-**SP15**-DVT (-Met1) | 14760.97 | 14760.7 |
| GB1-**SP15**-$(DVT)_2$ (-Met1) | 14981.2 | 14981.2 |

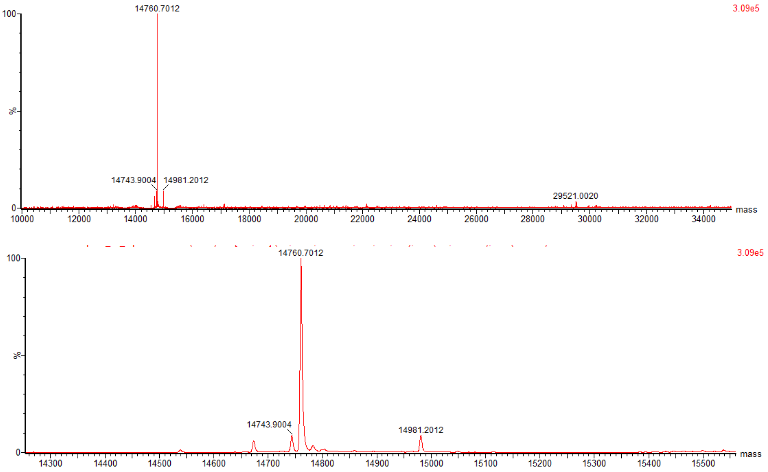

**Figure S15**. MS analysis of the GB1-fused peptide **SP15.**

### Mass spectra of free stapled peptides

## SP2

| Species | m/z calculated | m/z found |
|---|---|---|
| M+2H | 2289.97 | 2290.7 |
| M+3H | 1526.98 | 1527.5 |
| M+4H | 1145.48 | 1146.0 |
| M+5H | 916.59 | 917.0 |

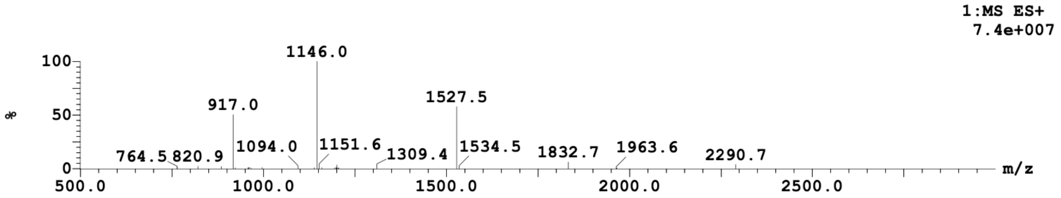

Figure S16. MS analysis of purified stapled peptide **SP2**.

## SP24

| Species | m/z calculated | m/z found |
|---|---|---|
| M+2H | 1483.72 | 1483.3 |
| M+3H | 989.48 | 989.3 |
| M+4H | 742.36 | 742.0 |
| M+5H | 594.095276 | 594.0 |

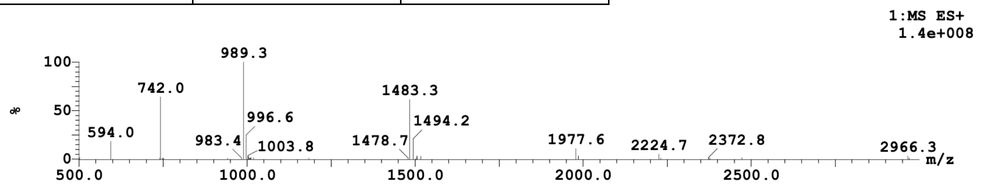

Figure S17. MS analysis of purified stapled peptide **SP24.**

## SP30

| Species | m/z calculated | m/z found |
|---|---|---|
| M+2H | 1527.267276 | 1526.9 |
| M+3H | 1018.513943 | 1018.2 |
| M+4H | 764.137276 | 764.0 |
| M+5H | 611.511276 | 611.5 |

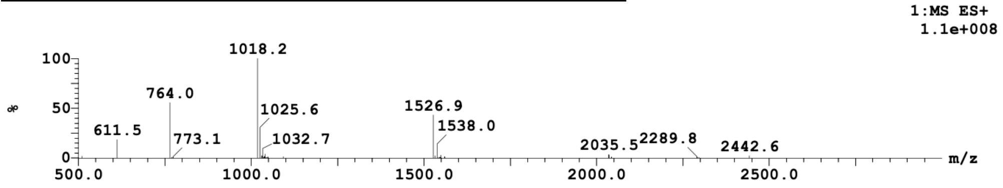

**Figure S18**. MS analysis of purified stapled peptide **SP30**.

## SP31

| Species | m/z calculated | m/z found |
|---|---|---|
| M+4H | 1144.08 | 1144.0 |
| M+5H | 915.46 | 915.3 |
| M+6H | 764.58 | 762.9 |
| M+7H | 654.84 | 654.1 |

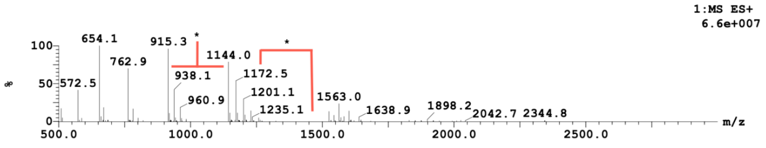

* = TFA counterions

**Figure S19**. MS analysis of purified stapled peptide **SP31**.

### HPLC analysis of free stapled peptides

## SP2

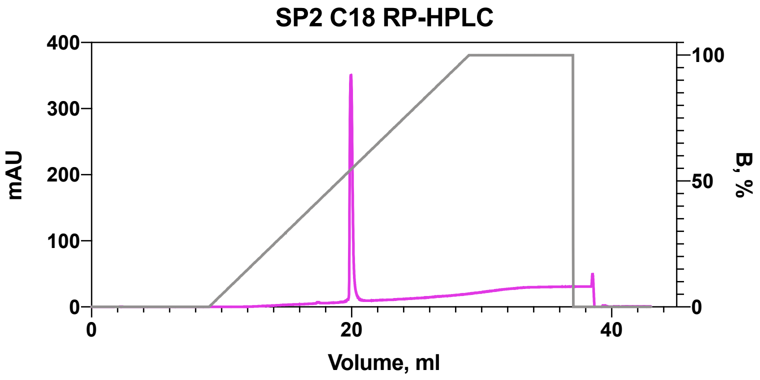

**Figure S20**. HPLC chromatogram of purified stapled peptide SP2.

## SP24

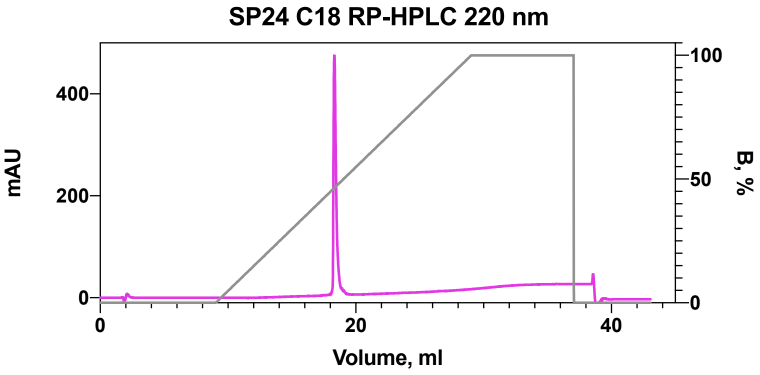

**Figure S21**. HPLC chromatogram of purified stapled peptide **SP24**.

## SP30

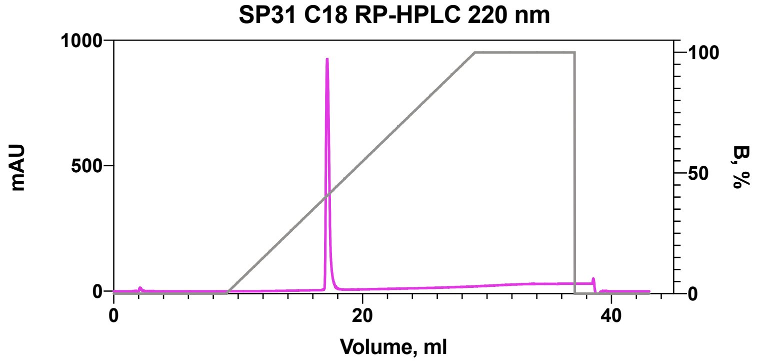

**Figure S22**. HPLC chromatogram of purified stapled peptide **SP30**.

**SP31**

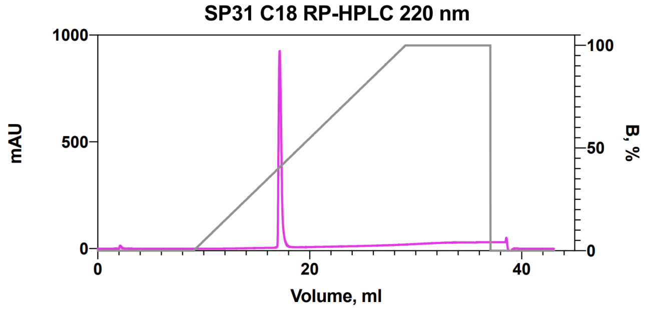

**Figure S23**. HPLC chromatogram of purified stapled peptide **SP31**.
